# Learning neural dynamics through instructive signals

**DOI:** 10.1101/2025.08.30.673300

**Authors:** Rich Pang, Juncal Arbelaiz, Jonathan W. Pillow

## Abstract

Rapid learning is essential for flexible behavior, but its basis in the brain remains unknown. Here we introduce the PRISM plasticity rule, a unifying mechanistic model of three well-established, fast-acting synaptic plasticity rules—in hippocampus, cerebellum and mushroom body—which relies exclusively on pre-synaptic activity and an “instructive signal” from another brain area. Using a multi-region network model we show that guiding PRISM plasticity with instructive signals enables the network to quickly learn extremely flexible nonlinear dynamics underlying behaviorally relevant computations, as well as to emulate unknown external system dynamics from real-time error signals, which we demonstrate with comprehensive simulations supported by exact mathematical theory. Thus, PRISM plasticity guided by instructive signals is well-suited to rapidly learn general-purpose neural computations—in contrast to canonical Hebbian rules. Finally, we show how including this plasticity rule in artificial learning algorithms can solve long-range temporal credit assignment, a long-standing challenge in machine learning.

**Highlights:**

- PRISM (PResynaptic and Instructive Signal-Mediated) plasticity—a unifying mechanistic model of three fast-acting plasticity rules in hippocampus, cerebellum, and mushroom body.
- Mathematical theory inspired by Support Vector Machine exactly predicts the dynamics learned in a network model through PRISM plasticity.
- Examples of learning nonlinear dynamics in a single shot via sparse instructive signals or from real-time error signals.
- Machine-learning application of instructive signals and PRISM plasticity to solve the long-range temporal credit assignment problem.

## INTRODUCTION

Rapid learning is essential for flexible behavior, allowing animals to quickly adapt to changing environments and task conditions. Mechanistically, learning is thought to be mediated by synaptic plasticity in the brain. To date, most models of biologically plausible learning focus on Hebbian plasticity, in which synaptic weight changes are driven by coincident pre- and post-synaptic activity^1–10^. Such rules have wide support from *in vitro* experiments ^3,4,11,12^ and in network models are well suited to form networks capable of performing pattern completion supporting memory recall, or the production of stereotyped activity sequences supporting motor outputs ^2,5,6,9^. It has proved difficult, however, to coerce Hebbian rules to produce more general-purpose computations needed for flexible behavior^13^, challenging their overall power and expressivity. Moreover, recent experiments suggest that some of the strongest plasticity rules *in vivo*, with timescales best matched to rapid learning (∼seconds), are in fact markedly non-Hebbian in character^14–18^. Thus, it sensible to consider what types of computations fast, non-Hebbian rules can produce.

In mammalian hippocampus and cerebellum, and the fly mushroom body—three highly plastic circuits involved in rapid learning—some of the strongest, most well-established synaptic plasticity rules depend not on pre- and post-synaptic activity, but on pre-synaptic activity and an “instructive signal” from another brain area^15–25^ (Fig 1a-b). We term this mechanism PRISM (“PResynaptic and Instructive Signal-Mediated”) plasticity. While the precise temporal dependencies vary^15,16,23^, PRISM plasticity in these three areas is highly stereotyped in several key aspects. First, it can act very quickly, producing large, lasting weight changes in just seconds. Second, it is largely independent of the post-synaptic neuron’s activity, in contrast to Hebbian plasticity. Third, it occurs at feed-forward output synapses from neural populations thought to exhibit *pattern-separated representations* (PSRs)—high-dimensional non-overlapping neural codes for upstream inputs^26–31^. PRISM plasticity at PSR outputs may thus reflect a common mechanism of rapid learning in the brain, yet its ability to produce flexible neural computations has received far less investigation than Hebbian rules.

**Figure 1:**
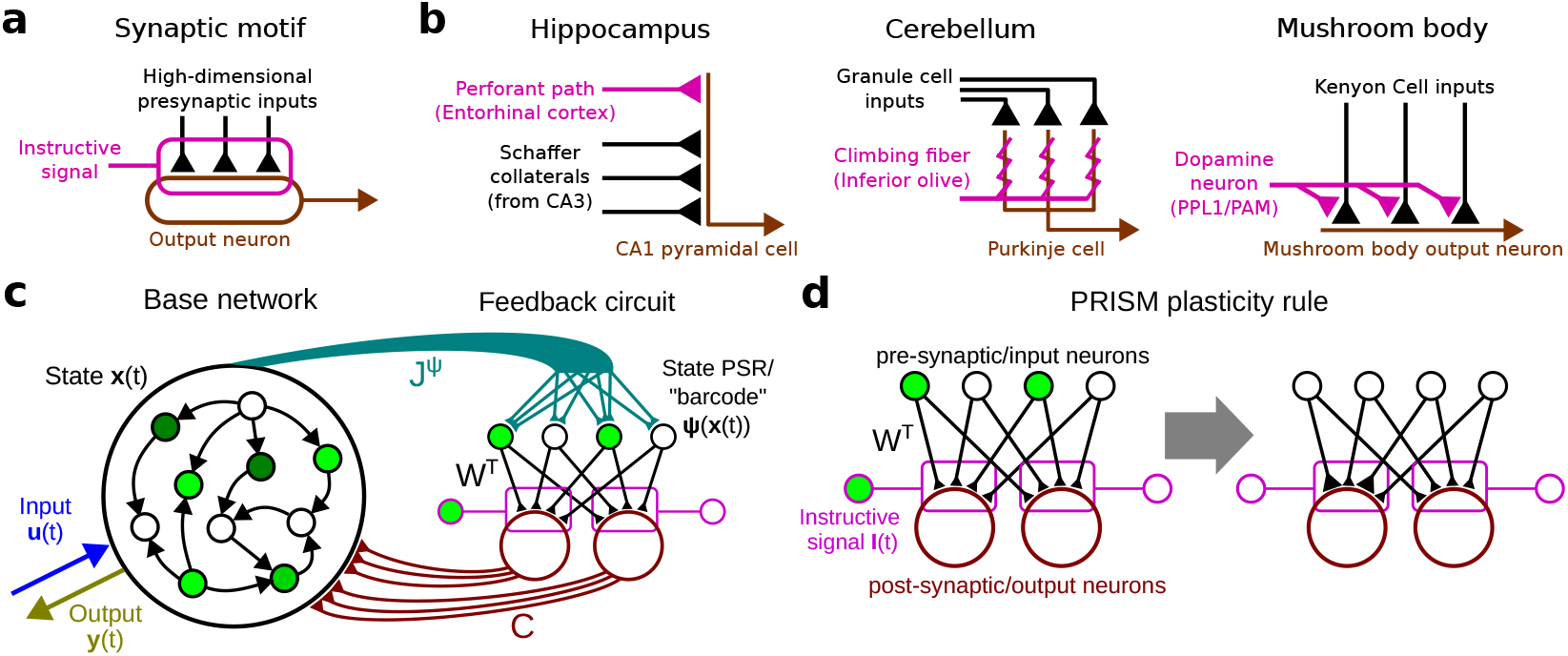
Network model and PRISM rule. **a**. Stereotyped synaptic motif in which plasticity of pre-synaptic weights onto post-synaptic output neuron depends on pre-synaptic activity and an instructive signal. **b**. Schematics of physical instantiations of synaptic motif in **a** in three different brain areas^14–18,20-23^. **c**. Network model including base network and plastic feedback circuit, with the latter shaped by instructive signals. **d**. Schematic of PRISM plasticity rule in Eq. ^(3).^

Computation in the brain is thought to be mediated by the activity dynamics of populations of neurons, which are often studied via recurrent neural network (RNN) models ^2,6,32-38^.An RNN is a flexible, nonlinear model that consists of a network of artificial neurons whose activities change over time under the influence of external and recurrent/feedback inputs. By adjusting synaptic weights, RNNs can be trained to perform diverse tasks and their internal activity subsequently examined to probe the learned dynamical mechanisms. Typically, RNN synaptic weights are learned through either back-propagation through time (BPTT)^37–39^, a flexible machine-learning algorithm yet usually considered non-biological; or Hebbian plasticity^2,6,8^, which has limited expressive power and may act too gradually in the brain to support rapid learning ^4,11,40,41^. While emerging work suggests that PRISM rules can enable RNNs to learn certain bespoke tasks^42,43^, whether they can produce dynamics supporting general-purpose nonlinear computation needed for flexible behavior is unknown.

Here we investigate a multi-region RNN consisting of a “base” RNN loosely analogous to neocortex and a feedback circuit analogous to hippocampus, cerebellum, or mushroom body. Synaptic outputs of PSR neurons within the feedback circuit are subject to PRISM plasticity guided by instructive signals external to the RNN. We show that given suitable instructive signals, PRISM plasticity enables the RNN to learn highly flexible, general-purpose nonlinear dynamics supporting behaviorally relevant computations, which we demonstrate with comprehensive simulations supported by exact mathematical theory. Our theory is precisely analogous to the well-understood theory of Support Vector Machines^44–46^, except set in a dynamical context, concisely linking the learned network connectivity to the resulting neural dynamics^34,35,47^. This in turn reveals the ability of PRISM rules to produce arbitrarily complex dynamics. Finally, we show how incorporating PRISM rules into artificial learning algorithms based on gradient descent can provide a solution for linking network outputs to long-vanished causal inputs, a long-standing challenge in machine learning^48^.

## RESULTS

We first introduce our network model and a mathematical theory for the rapid learning of neural dynamics. We then present a series of simulations characterizing several distinct ways the network can be trained through instructive signals, revealing the wide-ranging capabilities of rapid biological plasticity rules observed in the brain.

### Model

#### Architecture

Consider a “base” RNN reciprocally connected to a feedback circuit analogous to hippocampus (specifically, the CA3-to-CA1 layer), cerebellum, or mushroom body (Fig 1c). Let there be D neurons in the base RNN, and within the feedback circuit let there be N pre-synaptic or input neurons and Q post-synaptic or output neurons (Fig 1d). The pre-synaptic or input neurons correspond to CA3 pyramidal cells (hippocampus), granule cells (cerebellum), or Kenyon cells (mushroom body); and the post-synaptic or output neurons to CA1 pyramidal cells (hippocampus), Purkinje cells (cerebellum), or mushroom body output neurons. As in cerebellum and mushroom body, we will generally assume N≫Q, i.e. there is strong convergence from the pre-synaptic/input neurons to the post-synaptic/output neurons within the feedback circuit.

#### Dynamics

The activity dynamics of the neurons in the base RNN consist of four terms: a leak term reflecting the intrinsic decay of activity within each neuron, a term corresponding to recurrent inputs arising from within the base RNN, a term corresponding to recurrent inputs from the feedback circuit, and a term corresponding to external inputs. Specifically, let the dynamics of the *D* neurons in the base RNN follow:

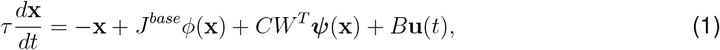

where **x***(t)* are the activity levels of the D base neurons, *ϕ* is an element-wise nonlinearity such as tanh, ***ψ***(**x**) are the PSR-like activities of the N input neurons within the feedback circuit (described in detail below), **u***(t)* are external inputs, and *τ* is a time constant determining the leak rate when all other inputs are zero. The recurrent synaptic weights within the base network are given by *J*^*base*^, the synaptic weights within the feedback circuit by *W*, the weights from the feed-back circuit output neurons back to the base RNN by *C*, and the weights on the external inputs by B. We optionally include a linear readout **y***(t)*. To focus on the effects of the feedback circuit we will usually consider *J*^*base*^ = 0, i.e. no base dynamics besides the linear decay toward the origin.

The input neuron activations ***ψ***(**x**) within the feedback circuit—corresponding to CA3, granule, or Kenyon cell activities—exhibit a PSR of the base RNN state **x**, created via a random projection of **x** by matrix *J*^*ψ*^ into an N-dimensional space followed by a nonlinearity. Unless otherwise noted, the main example we will use throughout this work, which we term the “Rand-Sig” ***ψ*** (random projection followed by a sigmoid), is:

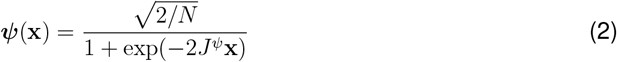

where the weights 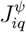 are sampled from a normal distribution with standard deviation, or “gain” *g*. The activities of the output neurons of the feedback circuit are given by **o**(*t*) = *W* ^*T*^ ***ψ***(***x***(*t*)) and are implicit in Eq. (1). Intuitively, the PSR ***ψ***(**x**) acts as a “barcode” for the system state **x**—each **x** maps to a unique (with high probability) high-dimensional vector ***ψ***(**x**), with largely uncorrelated barcodes for distant states but similar barcodes for nearby states, in line with the well-studied pattern-separation action of biological random networks^29,49^. As we will show, the form of ***ψ*** influences the geometric structure of the learned dynamics. For instance, surrounding Eq. (2) with a K-winner-take-all nonlinearity that silences all but the *K* most active neurons, as proposed in mushroom body and CA1 models^50,51^, can decrease interference and promote further pattern separation. However, we will usually consider the Rand-Sig ***ψ***.

#### PRISM Plasticity Rule

To focus on rapid plasticity mediated by instructive signals, we take inspiration from biological PRISM plasticity at PSR outputs and let all plasticity be localized exclusively to the weights *W* within the feedback circuit (Fig 1c). These weights are analogous to the feed-forward, convergent CA3 to CA1 projections (hippocampus), granule cell to Purkinje cell projections (cerebellum), and Kenyon cell to mushroom body output neuron projections. Our goal is to understand whether and how plasticity of these weights alone can produce flexible, general-purpose nonlinear dynamics, which would confer substantial power and expressivity to a single layer of feed-forward network weights subjected to a biological plasticity rule.

We formalize the PRISM rule as:

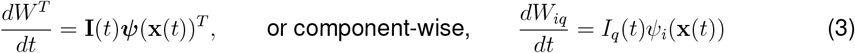

where **I**(*t*) is an external *instructive signal* with Q components, i.e. one instructive signal per output neuron. As in biology, PRISM plasticity is induced by pre-synaptic and instructive signals but expressed at synapses from pre-synaptic to post-synaptic neurons (Fig 1d). Note that PRISM, while non-Hebbian, is still a “two-factor” rule, in contrast to “three-factor” rules in which Hebbian plasticity is gated by an instructive or neuromodulatory signal^52–54^. For simplicity, Eq. (3) relies on simultaneous activation of the pre-synaptic neuron and the instructive signal, but see Extended Methods for a generalization to temporally extended plasticity windows^15–17,23^. The simultaneous case provides crucial mathematical insights for addressing the temporally extended case.

An illustration of the PRISM rule is shown in Fig 1d. A synapse *W*_*iq*_ from pre-synaptic neuron *i* to post-synaptic neuron *q* is modified when: (1) its pre-synaptic activity *ψ*_*i*_(***x***(*t*)) is greater than 0 (non-negativity is enforced by Eq (2)); and (2) there is a simultaneous instructive signal *I*_*q*_(*t) = 0*. If the instructive signal is positive (negative) the weight increases (decreases). The effect of the learning rule can also be understood as follows. At time *t*^*\**^, the barcode of system state ***x***(*t*^*\**^) is given by ***ψ***(**x**(*t*^*\**^)). If an instructive signal impulse *I*_*q*_(*t*)= *α*^*\**^*δ(t* − *t*^*\**^) occurs at this time, then column *q* of the weight matrix *W*, corresponding to the *q*-th output neuron, is incremented by *α*^*\**^***ψ***(**x**(*t*^*\**^)). Intuitively, ***ψ*** specifies “what” to learn, and **I** specifies “when” to learn it.

As we will show, different patterns of instructive signals **I**(*t*), along with external inputs **u**(*t*), can be used to train the network in different ways. In specific cases it will be natural to allow **I**(*t*) to follow an error signal between a target and the RNN trajectory, thus providing a “closed-loop” learning rule. Most generally, however, we remain agnostic as to the meaning or information content of **I**(*t*), and focus on its effects on connectivity and dynamics produced via PRISM plasticity.

### Theory of the learned flow fields produced through instructive signals

Flow fields are visual representations of a dynamical system depicting the magnitude and direction of forces *d****x****/dt* at different states, and which can be used to understand the system geometrically^55,56^. How do the instructive signals **I**(*t*), which update the weights *W* through the PRISM rule of Eq. (3), in turn update the flow field described by Eq. (1)? Linking RNN connectivity to dynamics and flow fields is a longstanding challenge in neuroscience^35,57–59^, and at first glance it may not seem that considering only connectivity and dynamics produced through PRISM plasticity would simplify this problem. Here, however, we show that the flow fields created through the PRISM rule can be expressed in a concise mathematical form precisely analogous to the “dual form” of Support Vector Machines—a widely studied and well-understood nonlinear machine-learning model—revealing a clear geometric interpretation of the learned dynamics.

#### Mathematical form of the learned dynamics

For simplicity, let the initial weights in the feedback circuit be zero (*W* = 0), and suppose the system is driven by inputs **u**(*t*), e.g. sensory stimuli, and subject to instructive signals **I**(*t*). We first consider point-process instructive signals expressed as a train of weighted impulses or delta functions,

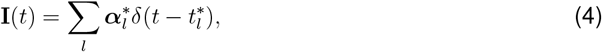

Where 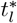 is the time of the *l*-th instructive signal impulse, and 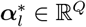 is a vector quantity giving the strength of each instructive signal component at the time of the impulse. We assume that all instructive signals are presented to the network during a bounded *instruction period*, after which **I**(*t*) is set to 0. Time-integrating the PRISM rule (Eq. (3)) and substituting the resulting weight matrix W into Eq. (1) reveals (see Extended Methods for derivation) that the learned dynamics following the instruction period can be written as

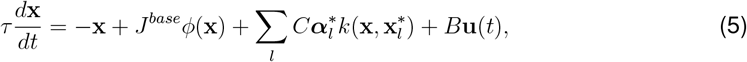

where the weight matrix *W* no longer appears and has been replaced by a summation over the instructive signals. The quantity 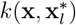 is a *kernel* function^46^ specifying the effective similarity between state **x** and state 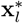 via the overlap of their barcodes, ***ψ***(**x**)^*T*^ ***ψ***(**x**^*\**^). Note that the final learned dynamics depend only implicitly, and not explicitly, on the timecourse of **u**(*t*) during the instruction period, which we discuss shortly. Eq. (5) is directly analogous to the “dual form” of SVMs, in which test outputs or predictions are expressed not in terms of weights, but in terms of how similar or near a test input is to a set of training inputs called *support vectors* ^46^. In our model, the flow field *d***x**/d*t* at an arbitrary state **x** is expressed in terms of how similar or near **x** is to a set of *support states* 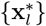. These support states are exactly the states of the base RNN when the instructive signals occurred: 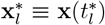. In analogy to SVMs, we term Eq. (5) the *dual form* of the learned dynamics. This dual form precisely links the learned high-dimensional network connectivity *W* to the resulting *D*-dimensional dynamics, which can be understood geometrically.

#### Geometric consequence of adding support states

The effect of an instructive signal impulse adding a support state can be understood geometrically by visualizing the corresponding change in the flow field. If we assume *J*^*base*^ = 0 and *W* = 0 initially, then prior to the instruction period the flow of the dynamics linearly draws all states back to the origin, due to the leak term − **x** in Eq. (5) (Fig 2a). Now drive the system with a time-varying input **u**(*t*) during the instruction period, which will move around the base RNN state. Applying a single instructive signal impulse at time 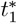 during this period adds a single support state 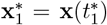 (Fig 2b). In accordance with Eq. (5) the resulting flow field becomes the sum of the base flow field −**x**/*ψ* and the patch of flow field, 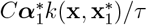, contributed by 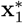. In the example of Fig 2c-d, the support state creates a second fixed point, revealing the capacity of PRISM plasticity to produce nonlinear dynamics, even when the base dynamics are fully linear. Thus, instructive signals add support states through the PRISM rule (Fig 2a-d), which concisely, yet precisely, describe the influence of the updated feedback circuit on the system dynamics.

**Figure 2:**
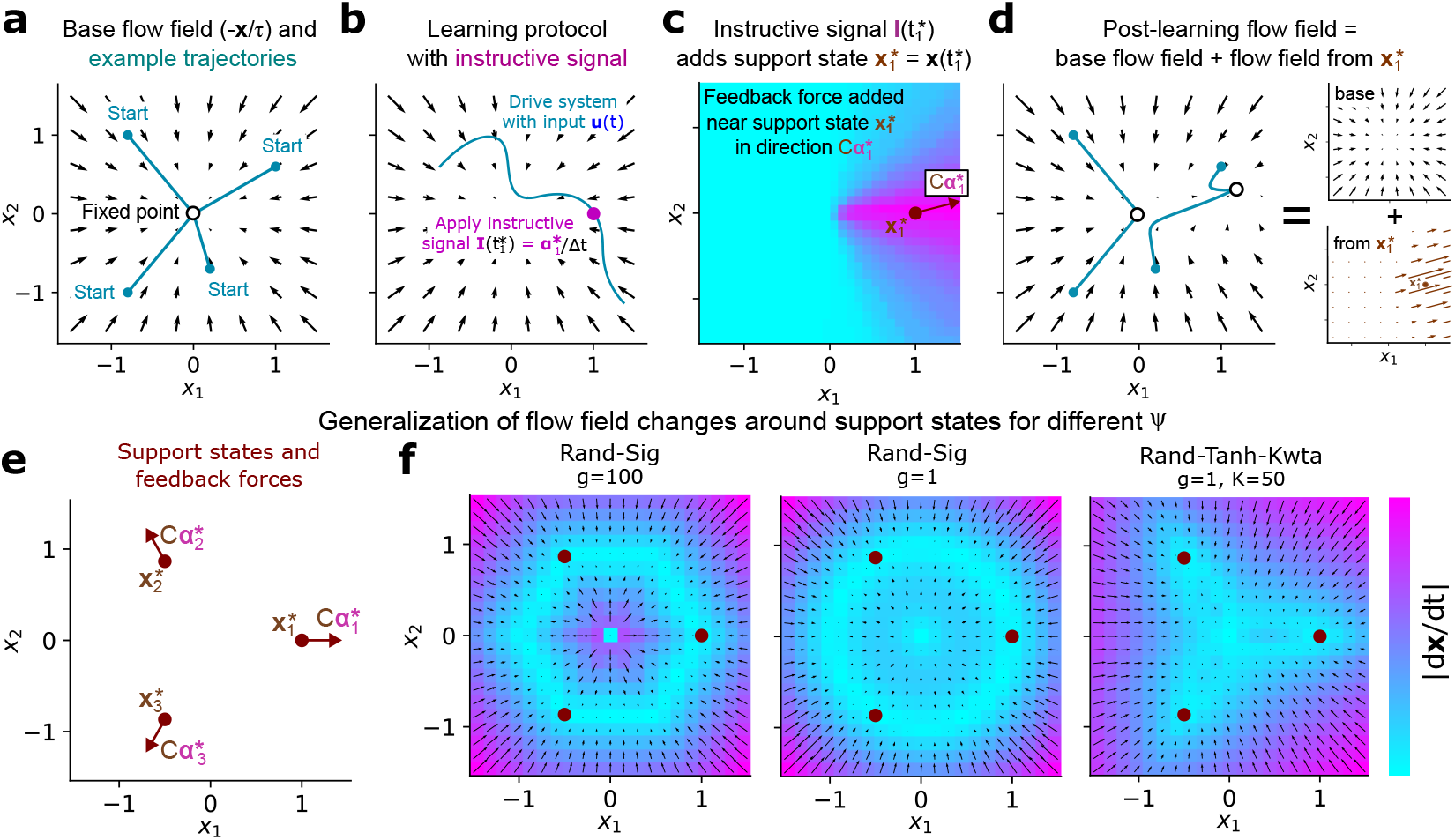
Flow fields produced by instructive signals. **a**. Example base flow field in which activity decays through uncoupled linear dynamics to a stable fixed point at the origin. **b**. Schematic of applying a single instructive signal impulse while the system is being driven by external input **u**(*t*). **c**. Schematic of the effect of the instructive signal in **b**, which is to add a support state at 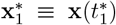, whose influence on the learned dynamics spreads according to 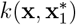. **d**. Schematic of final flow field after instruction period, which is a sum of the base flow field and the flow field contributed by the support state 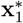. **e**. Schematic of three support states added to a 2-D state space with the same base dynamics as **a. f**. Flow field produced by the three support states in **e** for three different choices of ***ψ***.

Note that to compute the final flow field it does not explicitly matter *when* the instructive signal was applied or what inputs were presented during the instruction period—the only information retained explicitly in the learned flow field equation is *where* the system was in state space at 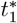 and the (vector) value 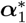 of the instructive signal. Multiple instructive signal impulses add multiple support states, each contributing its own patch of flow field around the support state, which is added to the sum in Eq. (5). (This statement holds for the PRISM rule in Eq. (3), but see Extended Methods for the generalization to temporally extended plasticity windows.) For continuous, rather than point-process instructive signals, the sum in (5) is replaced by an integral and the *set* of support states by a support state *density* (Extended Methods).

#### Generalization of the learned dynamics beyond the support states

Crucially, the manner in which the learned dynamics generalize to states not visited during the instruction period, or in which no instructive signal was applied, is determined by the kernel *k*, which defines the effective similarity between two states and in turn how a support state influences the dynamics at an arbitrary state. For instance, three support states optimized to produce three specific fixed points create different patterns of surrounding fixed and slow points when different ***ψ***(**x**) are used, since these yield different kernels (Fig 2e-f). Thus, how the learned flow field generalizes across the total state space can be controlled or predicted in a precise, straightforward manner through the choice of ***ψ***. Certain kernels, e.g. for the N, g → ∞ limit of the Rand-Sig nonlinearity (Eq. (2)), additionally admit a closed form that does not depend on *J*^*ψ*^ (Extended Methods), such that the dual form (Eq. (5)) depends on neither W nor *J*^*ψ*^. In this case, Eq. (5) reveals a concise, exact mathematical relationship between network connectivity of the complete RNN and the corresponding *D*-dimensional flow fields.

Our theory enables to precisely predict how sequences of instructive signals will update the dynamics flow field through the PRISM rule, even before we run the simulation forward, as long as we know the system state at the time of each instructive signal. Using this theory, we now address which flow fields can be produced through instructive signals **I**(*t*), and how the latter should be chosen to learn specific target dynamics.

### One-shot emulation of known target dynamics

Which dynamical systems can be expressed in terms of support states, and what patterns of instructive signals **I**(*t*) and external inputs **u**(*t*) can teach the RNN to produce these dynamics? We hypothesized that, unlike Hebbian plasticity—which produces dynamics largely limited to specific fixed-point attractors and sequences ^2,6,9,60^—the flexibility of instructive signal timing combined with the observation that each instructive signal impulse adds a new patch of flow field would allow our model to produce a wide variety of nonlinear flow fields. In this section, we first show that a large family of flow fields can be expressed in terms of support states, and then that these support states can be added online via suitable inputs and instructive signals.

To test our hypothesis, we considered the *N, g* → ∞ limit of the Rand-Sig nonlinearity (Eq. (2)), which we term the Rand-Sig-nonlinearity, and whose associated kernel function can be concisely expressed (Extended Methods) as

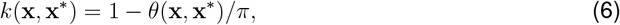

where *θ* is the *angle* between **x** and **x**^*\**^. Combining Eq. (5) and (6), the corresponding dual form for the dynamics is:

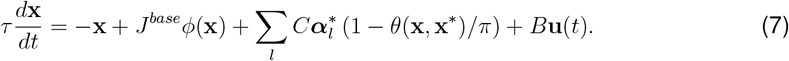

According to this equation, for the Rand-Sig-∞ nonlinearity, the learned component of the flow field does not depend on the magnitudes of **x** and **x**^*\**^, but only on *θ*. Hence, this learning rule and nonlinearity cannot produce changes to the flow field that vary with distance from the origin (however, the *total* flow field can still vary with distance to the origin due to the base dynamics). We verified this using ridge regression to identify loadings 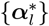 to produce target flow fields that did and did not satisfy this criterion (Fig S1). However, this restriction can be lifted by adding random biases in the nonlinearity ***ψ*** (Fig S1).

Even with this restriction, the flow fields that can be constructed in terms of support states with the Rand-Sig-***ψ*** are highly flexible. This is because each term in the sum in Eq. (7) acts like a basis function over the unit sphere in *D* dimensions, which can be combined arbitrarily combined. When *C* is the identity, for instance, then for *D* = 2, 3, any function on the sphere that can be expressed as the sum of a constant and an odd function can be constructed from support states, with a similar result likely holding for *D* > 3 (Extended Methods). To illustrate this point, we used ridge regression to identify support states and associated 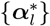 that can produce a variety of flow fields, including multiple fixed points, limit cycles, and an approximate line attractor, corroborating our mathematical result (Fig S2). Thus, support states can be combined to produce a wide variety of nonlinear dynamics thought to subserve neural computation^61–63^.

While the preceding ridge regression analyses show how flow fields can be decomposed in principle into support states, they do not show how these support states could be added online. We therefore asked whether such flow fields could be produced through appropriately patterned inputs **u**(*t*) and instructive signals **I**(*t*) via Eq. (3). If inputs and instructive signals exist that can add generic support states to the system, this would suggest that an animal could learn to “emulate” a target dynamical system in a brain area through sufficient **u**(*t*) and **I**(*t*). On the other hand, one might imagine that certain combinations of support states are unreachable through inputs and instructive signals.

We found that for identity *B* and *C*, and leveraging the radial invariance of the Rand-Sig-∞ kernel (Eq. (6)), we could construct a protocol for adding arbitrary support states through PRISM plasticity (Fig 3a). First, in an “offline” step (which we do not take to be biological) identify support states 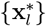 and loadings 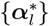 needed to produce a target flow field, e.g. using ridge regression. Second, perform the following: for each support state 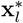 and loading 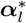, drive the system with a large input in the same direction as 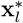, forcing the system into a state angularly close to 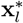 ; then apply an instructive signal impulse to trigger the PRISM rule and add the support state. Because support states at the same angular position are all equivalent (Eq. (7)), this provides a recipe for adding effectively arbitrary support states. Thus, any flow field expressible in terms of support states can emerge through appropriately timed inputs and instructive signals (Fig 3a).

**Figure 3:**
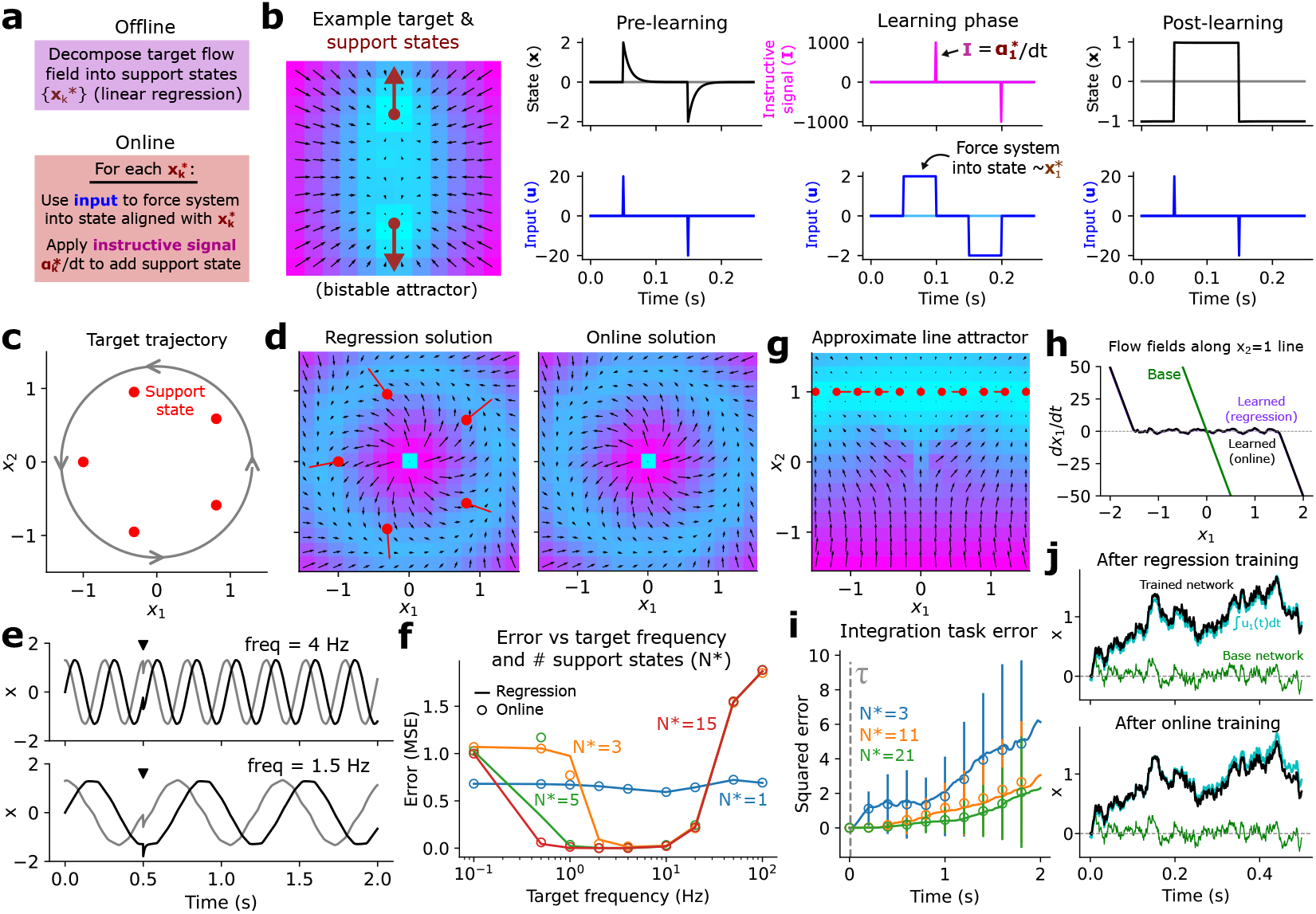
One-shot learning of known target dynamics. **a**. Offline-online algorithm for designing inputs **u**(*t*) and instructive signals **I**(*t*) to emulate a known target flow field. **b**. Example application of the algorithm in **a** to create a flow field with two fixed-point attractors. **c**. Schematic of flows/velocities to be learned along a specific trajectory by combining 5 support states. **d**. Left: regression solution (identifying 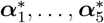) to approximate target flow in **c**. Right: solution produced by applying the online portion of the algorithm. **e**. Activity time-courses for **x**(*t*) when the system is trained to produce either of two example limit cycles. The black triangle shows the time of a small perturbation, indicating that the dynamics are stable with respect to this perturbation. **f**. Error between target and learned activity time-courses **x**(*t*) as a function of the target limit cycle frequency and the number of support states. **g**. Flow field and support states reproducing an approximate line attractor. **h**. Dynamics along the *x*_2_ = 1 line in **g** of the system trained to produce a line attractor that is used to integrate *u*_1_(*t*). The system is driven with an additional constant input *u*_2_(*t*) = 1 during this task. **i**. Error in using the trained system to perform integration along the line attractor, vs elapsed time and the number of support states afforded. **j**. Example of the trained network integrating its inputs.

We tested this procedure by training our network model to emulate three different example systems with known dynamics. First, we considered a target system with two stable fixed points. Fig 3b demonstrates an application of the offline-online algorithm (Fig 3a) to construct the inputs and instructive signals needed to produce this system given no base dynamics beyond linear decay to the origin. As predicted, the system learned the target dynamics after only two instructive signal impulses, allowing the network to subsequently act as a 1-bit memory.

Second, we asked whether the network could learn to produce a limit cycle at a specific target frequency. Of note, during the offline step, we can identify support states needed to produce only flows/velocities at specific states, rather than the entire flow field—for instance the flows along a circular trajectory (Fig 3c). We were curious, however, whether the learned dynamics around such a trajectory would be stable or unstable to perturbations, even if we taught the RNN to only reproduce velocities on the trajectory itself. We tested this by using the above procedure to teach the RNN to emulate the flow velocities along a circular trajectory with a target frequency *ν* (Fig 3c). As expected, we found that the trajectory velocities could be learned via inputs and instructive signals (Fig 3d). We also found, however, that the resulting dynamics were stable around the target trajectory, producing a limit cycle at the target frequency (Fig 3e). The range of frequencies of stable limit cycles increased with the number of support states or instructive signal impulses (Fig 3f). Thus, presenting inputs and instructive signals to produce a circular trajectory automatically stabilized the dynamics into a limit cycle across a range of frequencies.

As a third example, we asked whether our protocol could produce a line attractor—a canonical model of a neural dynamical motif supporting integration^33,61,64^. Indeed, our protocol could produce an approximate line attractor capable of integrating inputs over a timescale much longer than the leak timescale *ψ* (Fig 3g-j). Note, however, that for a finite number of support states the line attractor is not perfect. First, it is localized to a specific section of phase space, hence cannot integrate beyond its bounds (Fig 3h); second, it is composed of a microscopically rough landscape of slow and fixed points (Fig 3h). As a result, the inputs to be integrated need to be strong enough to drive the system around this landscape, but not so large as to escape the bounds of the attractor (Fig S3). Similar to the limit cycle example (Fig 3c-f), the stability of the line attractor against perturbations orthogonal to its longitudinal axis emerges automatically, even though we did not explicitly enforce this. Intuitively, this stability as well as that of the limit cycle arise from competition between the patches of flow field added by the instructive signals and the base flow field −**x**/*ψ*.

Thus, a rich variety of flow fields can be produced given proper inputs **u**(*t*) and instructive signals **I**(*t*). Moreover, this rule is highly efficient, requiring only sparse instructive signals—one impulse per support state to be added—yet which can qualitatively change the behavior of the RNN. Such a system is therefore well-posed for rapid or one-shot learning to emulate flexible dynamical systems, and may be highly resource-efficient if instructive signals are costly.

### Online learning of unknown dynamics through real-time error signals

We next asked whether our RNN could learn effectively through PRISM plasticity without having access to the target dynamics *a priori*. To address this we considered an online learning scheme in which a separate, reference system exists whose full dynamics are unknown but whose state **z**(*t*) can be observed during the instruction period. We asked whether an instructive signal could be derived from observations of the reference system state that would teach a learner RNN governed by Eq. (1)–(3) to eventually approximate the activity dynamics of the reference system without further instruction.

Leveraging tools from adaptive control theory (Extended Methods), we found that this was indeed the case. Suppose the true reference system follows dynamics

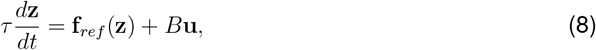

for some function **f**_*ref*_ unknown to the learner RNN. Let the learner RNN follow the dynamics

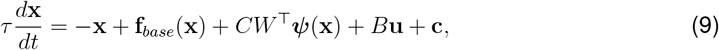

where **f**_*base*_ is e.g. *J*^*base*^*ϕ*(**x**), and which is equivalent to Eq. (1), except modified to explicitly include an additional input **c**, which we call a “corrective signal”. Specifically, let **c** ≡*k***e**, where **e**(*t*) ≡ **z**(*t*) **x**(*t*) is the tracking error between the reference and learner system state and *k* is a positive “control gain”. Let the weight updates and instructive signal follow

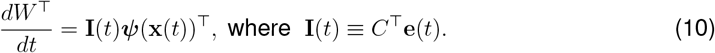

Note that the instructive signals here are continuous, rather than a point process. A schematic of the complete architecture is shown in Fig 4a.

**Figure 4:**
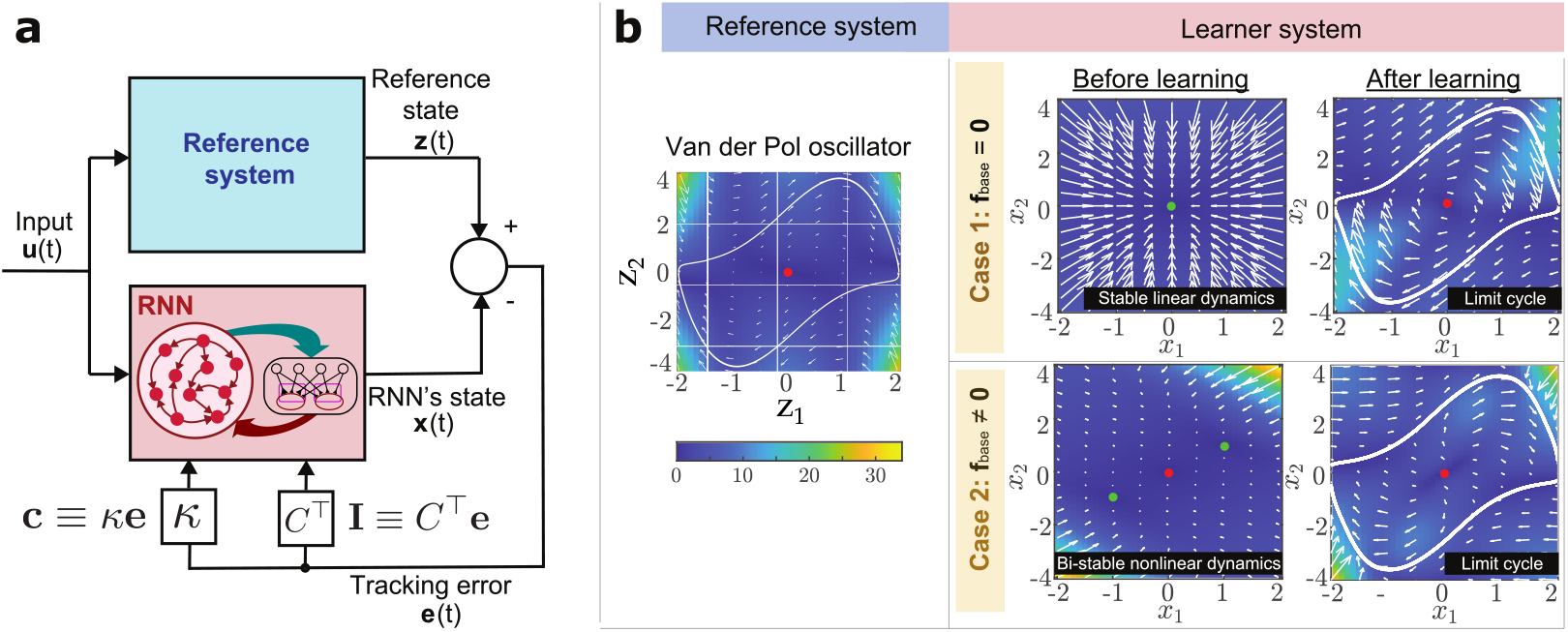
Learning to emulate an unknown system through real-time error signals. **a**. Architecture of learner RNN guided by error signals from a reference system. A more detailed block diagram is provided in Fig S4. The reference system and learner RNN both receive the same input B**u**. The difference between the reference’s and RNN’s states defines the tracking error **e**, which is used to generate the *corrective signal* **c** = *K***e** and the *instructive signal* **I** = *C*^T^**e** that are fed back into the RNN. **b**. Numerical case studies. The reference system is a Van der Pol oscillator, which has a stable limit cycle and an unstable fixed point at the origin. Case 1 has **f**_*base*_ **0** and exhibits the dynamics of a strictly stable linear system (due to the leakage term) before learning. In case 2, the base RNN dynamics support a bi-stable attractor. In both cases, the RNN successfully changes its local flow into stable limit cycle dynamics after the instruction period. Stable (unstable) fixed points are indicated by green (red) dots. Color-code of the flow fields is preserved across panels and indicated by the colorbar.

Our goal is for the learner RNN to learn to eventually reproduce the reference dynamics **f**_*ref*_ without requiring access to **z** or any further instruction. Thus, after the instruction period, we set **I** and *K* to zero, ceasing learning and letting the RNN behave independently of the reference system. The learning framework presented in this section is grounded on the theory of adaptive control^65,66^, specifically Model Reference Adaptive Control (MRAC). Under certain model matching assumptions and appropriate selection of the value of the gain *K*, convergence of the RNN’s state to that of the reference system can be asymptotically guaranteed using Barbalat’s lemma^67^ or through contraction theory^68–70^—methods used for the analysis of non-autonomous nonlinear dynamical systems. Further, selection of the inputs ***u*** to guarantee “persistency of excitation” conditions (intuitively, guaranteeing sufficient exploration of the state space) ensures learning of appropriate weight values^71,72^. Note that while the learner observes the reference system state, the reference receives no information from the learner. Such additional information exchange could make learning active and hasten learning rates; however, for simplicity we do not include it here, and it is not necessary for learning.

To validate the ability of the instructive signals (Eq. (10)) to teach the RNN to emulate the reference system, we numerically explored two illustrative case studies. In both cases, the reference dynamics to be emulated were described by a Van der Pol oscillator (Fig 4b, left), a 2-D nonlinear system that generates relaxation oscillations (Extended Methods). First, we analyzed the setting in which **f**_*base*_ ≡ **0** (i.e. linear uncoupled base dynamics decaying to the origin), while in the second case we consider pre-existing base dynamics supporting a bi-stable attractor, reflecting a scenario in which an existing learned flow field must be overwritten in order to emulate the limit cycle of the reference Van der Pol oscillator. We found that, regardless of the base dynamics considered, the RNN successfully learned a Van-der-Pol-like limit cycle resembling the reference system after the instruction period (Fig 4b, right). Thus, the corrective and instructive signals were sufficient to teach the network to emulate the reference dynamics through real-time feedback.

As here the instructive signals are continuous rather than a point process, the dual form of the learned dynamics after the instruction period requires a slight modification from Eq. (5). In particular, here the dual form of the dynamics is represented not in terms of individual support states, but via a support state *density* (Fig S7; Extended Methods). In other words, the sum in Eq. (5) is replaced by integral of the product of k with the support state density, once again revealing the flexibility of dynamics and flow fields that can be produced in this manner. Thus, through PRISM plasticity, instructive signals support both the online, one-shot implementation of known target dynamics (Fig 3), as well as the ability to learn online to emulate a reference system through real-time error signals (Fig 4). In both cases, the set of flow fields that can be produced by the learning rule is extremely flexible, due to the basis-function-like nature of the kernel function—the central limitation is the set of states that can be accessed during the instruction period, which determines where support states can be added.

We note the related idea of online internal model adaptation through error feedback studied in motor control. For example, it has been proposed that the cerebellum contains internal inverse and forward models of the motor apparatus^73^. The inverse models provide the neural commands necessary to achieve a desired motor trajectory or state and are interpreted as feedforward controllers. Forward internal models predict the behavior of the motor system in response to motor commands and can be leveraged, for instance, to compensate for time delays in feedback control^74^. Further, these models can in principle be learned within the cerebellum via different training signals: inverse models through motor command errors ^73,75,76^ and forward models through sensory prediction errors^73^. Thus, our results on instructive signals teaching the RNN to emulate reference dynamics are consistent with existing ideas in cerebellar motor control, while providing additional theoretical support and generality.

### Optimizing the instructive signals for long-range credit assignment

Finally, we asked whether instructive signals and PRISM plasticity could be leveraged to train RNNs on tasks involving long-range temporal credit assignment, which challenges traditional RNN training methods. Canonical RNN training in machine learning, typically based on back-propagation through time (BPTT), updates the RNN weights to minimize a loss function by descending the loss gradient^39,77^. However, while BPTT is both flexible and admits biological approximations^78,79^, training in the face of delayed loss evaluation is sensitive to “vanishing (or exploding) gradients”—these arise due to the repeated multiplication of the weight matrices across timesteps when calculating loss gradients^48,80^. In practice, this hinders temporal credit assignment between outputs that are separated from their causal inputs by long delays.

Here we give a proof of concept demonstrating how instructive signals can be used to address this problem by avoiding vanishing/exploding gradients. In our proposed method, which we term “instructor training”, we do not optimize the weights directly but instead optimize the instructive signal timecourse. For simplicity, here we consider the discretized version of our RNN, in which **x**(*t*), **I**(*t*) are replaced by **x**^*t*^, **I**^*t*^, etc. Given a fixed input sequence **u**^*t*^, each candidate instructive signal timecourse **I**^*t*^ produces a trajectory of the weight matrix W ^*t*^ according to the PRISM rule, and which after the instruction period yields a final, fixed W. We then optimize **I**^*t*^ using the same inputs and delayed loss function as one would in BPTT. Our final goal is to be able to run the initial plastic RNN forward (Eq. (1)–(3)) while applying the optimized instructive signals, such that the final weights W at the end of the instruction period allow the RNN to solve the task with no further instruction. Identifying task-optimized instructive signals could also provide a means to predict instructive signal dynamics in animals throughout task learning^81^.

Crucially, optimizing the instructive signal timecourse **I**^*t*^ can converge to at least two possible outcomes. First, the optimal instructive signals could remain strong throughout the course of learning, corresponding to a case in which the network learns to rely on the instructive signals to solve the task, thus never learning to perform the task independently. More favorably, the optimal instructive signals could decrease over learning, i.e. with the network initially relying on the instructive signals but by the end of the instruction period only relying on the weights W —in this case the network learns to solve the task independently without further instruction. We hypothesized that it was possible for the second case to emerge automatically simply by optimizing the instructive signal timecourse directly, even without imposing an explicit decay of their magnitude.

We tested our hypothesis on a proof-of-concept task with loss evaluated at a variable delay. That is, the loss depends only on the network activity occurring after the input has ceased (Fig 5a). To illustrate the essential challenge of long-range temporal credit assignment, we considered a parsimonious “hold” task with a delayed loss evaluation (Fig 5b). Here, in response to an input pulse in the [− 1, 0] direction, the network should respond by holding the first neuron’s activity x_1_ at -1, and vice versa for an input pulse in the [+1, 0] direction. Loss evaluation (the error between the true x_1_ and the target), however, only occurs after a variable delay following each input. As shown in Fig 5c-d, both BPTT^39,77^ and e-prop (a biological version of BPTT)^79^ succeed for short delays (on the order of the network time constant *ψ*) but fail for longer delays, as expected from the vanishing gradient problem^48^.

**Figure 5:**
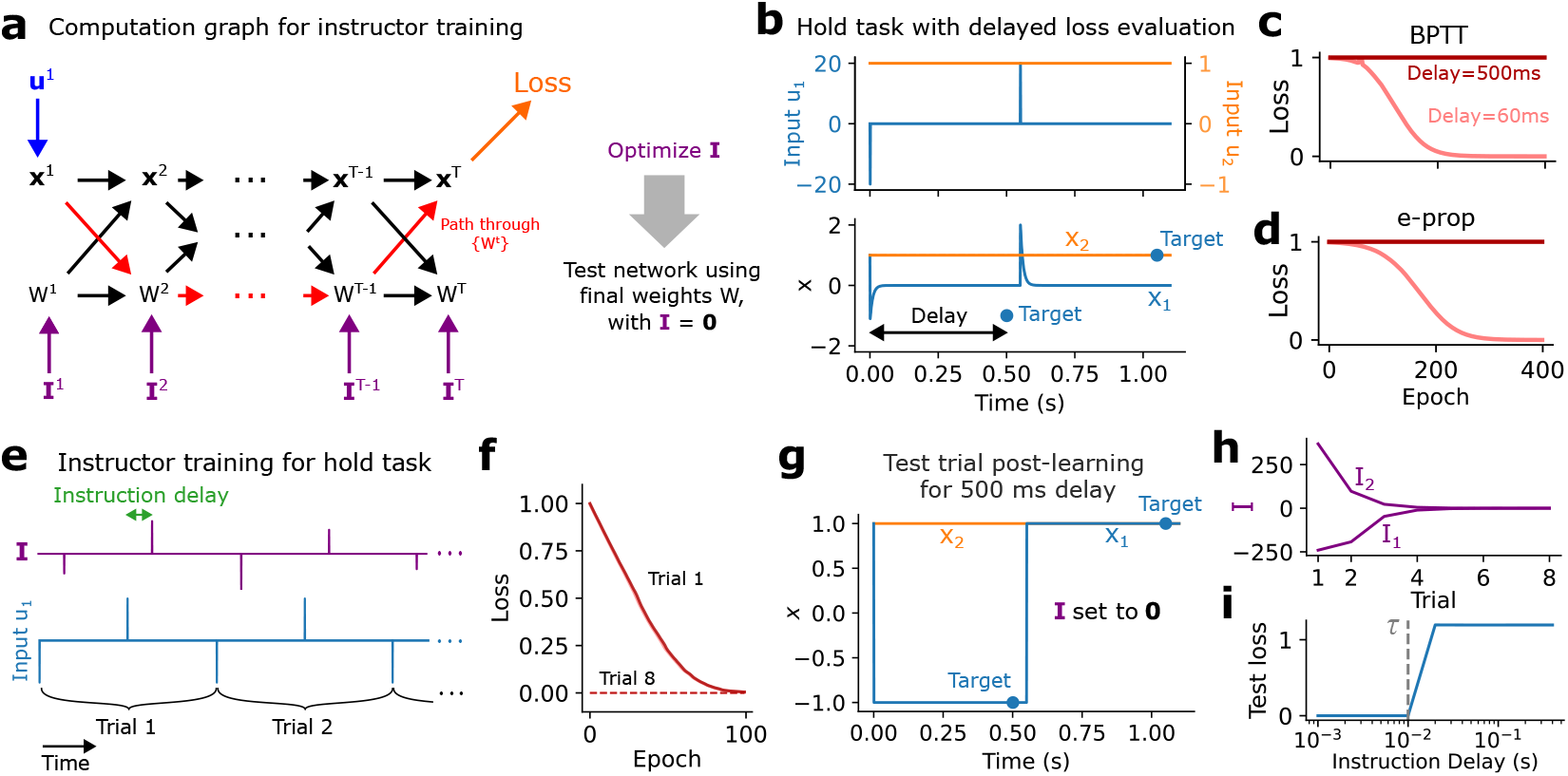
Instructor training proof-of-concept. **a**. Computation graph for instructor training. **b**. Example task in which loss is evaluated only after a delay, which is expected to trigger the vanishing gradient problem for BPTT when the delay is too long. The goal is for *x*_1_ (blue) to pass through each of the targets. **c, d**. Loss vs training epoch for the hold task in **b** for BPTT and e-prop for two different delays. **e**. Schematic of instructor training paradigm used for the hold task. Here the magnitudes of the instructive signal impulses are optimized. **f**. Loss vs epoch for each trial in the instructor training paradigm of **e**, given a 500 ms delay. **g**. Network performing hold task after instructor training, and with the instructive signal silenced. **h**. Magnitude of each of the two instructive signals vs trial number. **i**. Test loss (corresponding to the test trial in **g**) as a function of the instruction delay.

We found, however, that instructor training could solve this task, even with long delays, and moreover without learning to rely on the instructive signal by the end of training. Specifically, we considered 8 sequential “trials” of the hold task, each containing two inputs, with one instructive signal impulse allowed after each input (Fig 5e). This represents an approximate version of optimizing the complete instructive signal timecourse. We then optimized the amplitudes of the instructive signals for each trial, using the resulting weights W at the end of each trial as the initial weights for the subsequent trial. Indeed, for a long delay (500 ms, which neither BPTT nor e-prop can handle) the loss decreased over both training epochs within a trial and over trials. (Note also that grouping BPTT or e-prop training epochs into trial groups does not help them solve this task.) Importantly, in a test trial in which no instructive signals were included, the network still solved the task (Fig 5g), indicating that it had learned to rely on the weights rather than the instructive signals. Indeed, the magnitude of the optimized instructive signals decreased over trials (Fig 5h). Finally, we found that in order for the network to learn to solve the task without relying on the instructive signals it was necessary for the instructive signals to occur shortly after the inputs (Fig 5i).

Instructor training succeeds for the following reason. Formally, loss gradients needed for optimization via gradient descent are computed by summing over all paths through the computation graph from the network parameters to the loss (Fig 5a, S5), with each path corresponding to a product of Jacobians (the matrix of partial derivatives of the output with respect to the input variables) along that path^82^. In BPTT, however, the paths linking inputs to loss evaluation through the network states {**x**^*t*^} often “vanish” or “explode” if there is a long delay between the input and loss evaluation^48,80^, due to the repeated multiplication of the Jacobians in the gradient calculation. In instructor training, however, the weights are themselves dynamic variables, opening additional paths through the computation graph—i.e., through {W^*t*^} —for information from past inputs to reach the loss. Because the PRISM rule (Eq. (3)) has no decay time constant, this allows information to persist in the weights over long time periods, which in turn allows gradient information to propagate through the computation graph without vanishing or exploding. Intuitively, the optimization process learns that an instructive signal can be invoked to store information about the inputs in W until it is needed again; at the same time, however, it learns over sequential trials to transfer this information storage ability into the dynamics encoded in *W*.

Thus, instructor training represents an alternative to canonical gradient methods for end-to-end training of RNNs. In both BPTT and instructor training, the final result is an RNN with fixed weights *W*. Instructor training, however, leverages instructive signals to learn long-range dependencies during the training process, even though at test time no instructive signals or plasticity are required. Note that we do not treat the process of optimizing the instructive signals as biologically plausible; however, the optimal instructive signals reflect a biological solution to updating the weights if those instructive signals could be generated in the brain. These results thus suggest a biologically inspired solution to a long-standing problem in machine learning.

## DISCUSSION

We have shown how biologically inspired PRISM plasticity rules mediated by pre-synaptic activity and instructive signals can rapidly produce highly flexible nonlinear network dynamics. This work represents a complement to the theory of Hebbian plasticity’s ability to produce fixed-point attractors or sequences that can be used for pattern completion ^2,5,6,9^. Instructive signals, when applied to a plastic feedback circuit coupled to a base RNN, add support states to the dynamics flow field that can be combined to yield arbitrary nonlinear dynamics. A notable feature of the learning system we have proposed is that addition in the space of synaptic weights (via Eq. (3)) corresponds directly to addition in the flow field (Eq. (5))—thus, PRISM plasticity applied to feedback circuits reveals a simple, interpretable link between synaptic weights and flow fields, addressing the long-standing challenge of linking RNN connectivity and dynamics ^34,35,47,57^. As PRISM plasticity also accords more strongly with experimental data on rapid learning than Hebbian plasticity^15–17,23,42^, our model may additionally be more applicable to explaining empirical neural dynamics.

Our work extends a growing body of work on training RNNs with biological learning rules. One avenue of past research, encompassing the “RFLO”^78^, “e-prop”^79^, and MDGL^83^ algorithms, focused on how loss gradients can be approximated by local cellular processes in a single RNN. A second avenue of research has focused on dynamics and learning in RNNs coupled to a plastic feedback circuit analogous to a cerebellum^43^ or mushroom body^42^, or more generically an adaptive feedback controller^66,84^, similar to our model architecture (Fig 1c); however, while such RNNs can solve interesting tasks in a more biologically plausible manner than traditional training algorithms like BPTT, the mathematical relationship between the learning rule, the networks weights, and the resulting flow fields has remained elusive. Although some theoretical work has established important links between network connectivity and flow fields^35,85^, these studies have largely not considered connectivity produced by rapid biological plasticity rules. Our work synthesizes the above by providing a unifying framework and concise mathematical theory that links network connectivity produced through rapid biological learning rules to interpretable nonlinear dynamics and flow fields. Our study also suggests an RNN algorithm—instructor training—that can in certain cases solve long-range temporal credit assignment problems by overcoming the vanishing/exploding gradient problem, which hinders learning in canonical algorithms^48^ unless additional slow variables are introduced that can be used at test time^79,86^.

Our work represents a simplified model of how biological plasticity rules shape neural dynamics to support rapid learning and remains subject to important limitations. We have largely considered a plasticity rule in which weight changes are triggered by precise coincidence between presynaptic activity and the instructive signal, whereas empirical plasticity rules in hippocampus, cerebellum, and mushroom body exhibit various extended temporal windows of coincidence detection^15–17,23^. The dual form of the learned dynamics (Eq. (5)) can be generalized to accommodate such temporally extended windows (Fig S6, Extended Methods), but in future work it will be important to establish in detail how such modifications influence learning. Additionally, in all of our examples we have considered the case in which the (columns of the) input and feedback connections, *B* and *C*, respectively, span the entire space R^*D*^ of the base dynamics. In general, however, the columns of *B* and *C* might span only subspaces of R^*D*^, and it will be important to characterize which dynamics and tasks can be learned under these constraints. Finally, it will be useful to explore how different base dynamics **f**_*base*_ impair or accelerate learning, and to characterize in more detail the tasks that can be solved via instructor training.

Our theory makes testable general predictions, while also opening an avenue for fitting to data to make predictions in specific experimental contexts. The first general prediction is that activating an instructive signal will alter the neural dynamics in other brain areas predominantly around the system state these dynamics were in at the time of the instructive signal—the support state. This could be tested, for instance, by analyzing neural activity in cortical areas while stimulating climbing fibers inputs to cerebellum or perforant pathway inputs to hippocampus. Second, many different sequences of instructive signals can ultimately produce the same or similar flow fields—thus, for animals that all learn a task successfully, we predict there to be substantial heterogeneity in the instructive signals evoked throughout learning, even if the learned neural dynamics are similar across animals^87^. Finally, our model can in principle be fit to empirical neural population activity trajectories by identifying support states that would produce dynamics consistent with those trajectories. Such an approach could be used to make predictions about unseen neural responses in regions of state space not visited by the training data, benefiting from the inductive bias encoded by the kernel function.

## METHODS

Methods details and mathematical derivations are available in the Extended Methods section accompanying this manuscript.

## RESOURCE AVAILABILITY

### Lead contact

Requests for further information and resources should be directed to and will be fulfilled by the lead contact, Rich Pang.

### Data and code availability

- This project generated no new datasets.
- All code used to generate the figures in this manuscript is available at github.com/rkp8000/prism-rnn.
- Any additional information required to reanalyze the data reported in this paper is available from the lead contact upon request.

## ACKNOWLEDGMENTS

This work was funded by The Princeton Neuroscience Institute Innovation Award (to RP and JWP), The Princeton Neuroscience Institute C.V. Starr Award (to JA), the NIH BRAIN initiative (R01EB026946; to JWP), the NIMH (R01EY033064; to JWP), a U19 NIH-NINDS BRAIN Initiative Award (5U19NS104648; to JWP), and the Simons Collaboration on the Global Brain (SCGB AWD543027; to JWP). We thank Hakan Tureci for insightful discusions during the development of this project, as well as Anushri Arora, Victor Geadah, and other members of the Pillow lab for many helpful thoughts and conversations.

## AUTHOR CONTRIBUTIONS

Conceptualization, R.P., J.A., and J.P.; methodology, R.P., J.A., and J.P.; investigation, R.P. and J.A.; writing-–original draft, R.P. and J.A.; writing-–review & editing, R.P., J.A., and J.P.; funding acquisition, R.P., J.A., and J.P.; resources, R.P., J.A., and J.P.; supervision, J.P.

## DECLARATION OF INTERESTS

The authors declare no competing interests.

## SUPPLEMENTAL INFORMATION INDEX

Figures S1-S7 and their legends in a PDF

## Notes

https://github.com/rkp8000/prism-rnn

### Competing Interest Statement

The authors have declared no competing interest.

### Summary of Updates

The plasticity rule has been renamed to PRISM (PResynaptic and Instructive Signal Mediated) plasticity. Additionally, minor text edits have been introduced throughout the manuscript for clarity.

