## Extended Methods and Supplemental Figures for "Learning neural dynamics through instructive signals"

#### Derivation of the dual form for point-process instructive signals

For simplicity we first consider point-process instructive signals. Let the $q$-th instructive signal consist of a series of impulses at times $\{t^*\}$, weighted by $(\alpha_l^*)_q$ i.e.

$$\mathbf{I}(t) = \sum_l \boldsymbol{\alpha}_l^* \delta(t - t_l^*) \qquad \text{or component-wise} \qquad I_q(t) = \sum_l (\alpha_l^*)_q \delta(t - t_l^*). \tag{11}$$

Integrating Eq. (3), the change in $W_{:,q}$, the $q$-th column of $W$, after time $T$ is given by

$$\Delta W_{:,q} = \int_0^T I_q(t)\boldsymbol{\psi}(\mathbf{x}(t))dt = \int_0^T \left(\sum_l (\alpha_l^*)_q \delta(t - t_l^*)\right) \boldsymbol{\psi}(\mathbf{x}(t))dt = \sum_l (\alpha_l^*)_q \boldsymbol{\psi}(\mathbf{x}_l^*) \tag{12}$$

where $\mathbf{x}_l^* \equiv \mathbf{x}(t_l^*)$ are the support states added by the instructive signal. Thus, $\Delta W_{:,q}$ depends only on the system states visited at the time of the instructive signal impulses and moreover equals the weighted sum of $\boldsymbol{\psi}(\mathbf{x}_l^*)$ evaluated at these states. Generalizing Eq. (12) to the change in the full weight matrix $W$ produces

$$\Delta W^T = \sum_l \boldsymbol{\alpha}_l^* \boldsymbol{\psi}(\mathbf{x}_l^*)^T. \tag{13}$$

Plugging $W = W_0 + \Delta W$ back into Eq. (1), we arrive at

$$\tau \frac{d\mathbf{x}}{dt} = -\mathbf{x} + \mathbf{f}_{base}(\mathbf{x}) + C\left(W_0^T \boldsymbol{\psi}(\mathbf{x}) + \sum_l \boldsymbol{\alpha}_l^* k(\mathbf{x}, \mathbf{x}_l^*)\right) + B\mathbf{u}(t), \tag{14}$$

where $k(\mathbf{x}, \mathbf{x}^*) = \boldsymbol{\psi}(\mathbf{x})^T \boldsymbol{\psi}(\mathbf{x}^*)$ is the kernel function. However, if $W$ was created exclusively through the learning rule, i.e. $W_0 = 0$, then

$$\tau \frac{d\mathbf{x}}{dt} = -\mathbf{x} + \mathbf{f}_{base}(\mathbf{x}) + C\sum_l \boldsymbol{\alpha}_l^* k(\mathbf{x}, \mathbf{x}_l^*) + B\mathbf{u}(t). \tag{15}$$

This form of our dynamics highly resembles the "dual form" of prediction functions in SVMs, in which predictions are written not in terms of weights (the primal form) but in terms of a subset of the training data, called *support vectors*. Hence we call $\{\mathbf{x}^*\}$ *support states*.

Note that we can also write Eq. (12) as a *spatial* integral:

$$W_{:,q}^T \boldsymbol{\psi}(\mathbf{x}) = \int \rho_q(\mathbf{x}^*) k(\mathbf{x}, \mathbf{x}^*) d\mathbf{x}^* \quad \text{with} \quad \rho_q(\mathbf{x}^*) \equiv \sum_l (\alpha_l^*)_q \delta(\mathbf{x}^* - \mathbf{x}_l^*), \tag{16}$$

i.e. $\rho_q(\mathbf{x})$ is a set of delta functions at the support states $\{\mathbf{x}_l^*\}$ weighted by $\{(\alpha_l^*)_q\}$. This is because

$$\int \rho_q(\mathbf{x}^*) k(\mathbf{x}, \mathbf{x}^*) d\mathbf{x}^* = \int \sum_l (\alpha_l^*)_q \delta(\mathbf{x}^* - \mathbf{x}_l^*) k(\mathbf{x}, \mathbf{x}^*) d\mathbf{x}^* = \sum_l (\alpha_l^*)_q k(\mathbf{x}, \mathbf{x}_l^*). \tag{17}$$

The left-hand side is a more general form of the right-hand side. This is essentially the same relationship that a charge density has to a collection of point charges in electrostatics.

Crucially, although $(\alpha_q^*)_l$ corresponds to the instructive signal $I_q(t)$ at time $t_l^*$, the only thing that actually matters is *where* the system was at $t_l^*$. In other words, from Eq. (17), every time an instructive signal impulse arrives one can think of adding to $\rho_q(\mathbf{x}^*)$ the term $(\alpha_l^*)_q \delta(\mathbf{x}^* - \mathbf{x}_l^*)$.

While this indeed corresponds to the value of $I_q$ at time $t_l^*$, once $\rho_q$ is updated, one can forget about $t_l^*$, since now $\rho_q$ contains all the information needed to compute the post-instruction flow field. Note that it also doesn't matter how $\mathbf{I}(t)$ was constructed (e.g. if it was predetermined or based on some error signal).

#### Derivation of the dual form for continuous instructive signals

We can write down the continuous version of Eq. (11) as

$$\mathbf{I}(t) = \boldsymbol{\alpha}(t) = \int \boldsymbol{\alpha}(t^*)\delta(t - t^*)dt^* \qquad \text{or component-wise} \qquad I_q(t) = \int \alpha_q(t^*)\delta(t - t^*)dt^*. \quad (18)$$

Intuitively, one can think of this integral as the limit of the sum in Eq. (11) when the intervals between impulses go to zero. Following similar reasoning as above, the continuous version of Eq. (16) becomes

$$\rho_q(\mathbf{x}^*) \equiv \int \alpha_q(t^*)\delta(\mathbf{x}^* - \mathbf{x}(t^*))dt^* \quad (19)$$

where the discrete index $l$ has now been replaced with the "continuous index" $t^*$. Note again that after this integration $\rho_q(\mathbf{x}^*)$ depends only on $\mathbf{x}^*$.

We now arrive at Eq. (16) (above), the expression for $W_{:,q}^T\boldsymbol{\psi}(\mathbf{x})$ for the continuous case. This is exclusively an integral over $\mathbf{x}^*$ i.e. over the state space of the original system. The most general expression is

$$W^T\boldsymbol{\psi}(\mathbf{x}) = \int \boldsymbol{\rho}(\mathbf{x}^*)k(\mathbf{x}, \mathbf{x}^*)d\mathbf{x}^*, \quad (20)$$

where $\boldsymbol{\rho}(\mathbf{x}^*) \equiv [\rho_1(\mathbf{x}^*), \ldots \rho_Q(\mathbf{x}^*)]$. As before, the above holds for any $\mathbf{I}(t)$ or $\boldsymbol{\alpha}(t)$, regardless of whether they were pre-determined or followed e.g. an error signal.

Note that the relationship between Eq. (13) (point-process instructive signal and discrete support states) and Eq. (16) (continuous instructive signal and support state density) is exactly analogous to the relationship between a set of point charges and a charge density in electrostatics, where $k(\cdot, \cdot)$ plays the role of the Green's function $G(\cdot, \cdot)$, and the output of the sum/integral the electric field or voltage[88].

#### Linear decomposition of target flow field into support states

Here we use linear regression to identify the $\{\boldsymbol{\alpha}_l^*\}$ that should be associated with a set of support states in order to reproduce a known target flow field. Let

$$\mathbf{F}(\mathbf{x}) \equiv \frac{d\mathbf{x}}{dt} = \frac{1}{\tau}\left(-\mathbf{x} + C\sum_{l=1}^{N^*} \boldsymbol{\alpha}_l^* k(\mathbf{x}, \mathbf{x}_l^*)\right) \quad (21)$$

describe the velocity/flow field at position $\mathbf{x}$, with $\mathbf{F}(\mathbf{x}) \in \mathbb{R}^D$, i.e. the flow field created out of support states $\{\mathbf{x}_l^*\}$ with $\mathbf{f}_{base} = 0$. Given $\{\mathbf{x}_l^*\}$ we can use linear (Ridge) regression identify $\{\boldsymbol{\alpha}_l^*\}$ needed to approximate a target flow field $\mathbf{F}^{targ}$. Taking $C$ to be the identity for simplicity (such that $D = Q$, $d = q$), and rearranging terms in the $d$-th component of $\mathbf{F}$, we would like

$$f_d^{targ}(\mathbf{x}) \equiv \tau F_d^{targ}(\mathbf{x}) + x_d = \sum_l (\alpha_l^*)_d k(\mathbf{x}, \mathbf{x}_i^*) \quad (22)$$

Letting $\tilde{\mathbf{f}}_d^{targ} \equiv [f_d^{targ}(\tilde{\mathbf{x}}^1), \ldots, f_d^{targ}(\tilde{\mathbf{x}}^{\tilde{N}})]^T$ be $f_d^{targ}$ evaluated at a set of $\tilde{N}$ grid points, and similarly $\tilde{\boldsymbol{\alpha}}_d^* \equiv [\boldsymbol{\alpha}_d^*(\tilde{\mathbf{x}}^1), \ldots, \boldsymbol{\alpha}_d^*(\tilde{\mathbf{x}}^{\tilde{N}})]^T$ we can write

$$\mathbf{f}_d^{targ} = \tilde{K}\tilde{\boldsymbol{\alpha}}_d^*, \tag{23}$$

where $\tilde{K} \in \mathbb{R}^{\tilde{N}\times N^*}$ is the matrix of kernel values between each grid point $\tilde{\mathbf{x}}$ and each support state $\mathbf{x}^*$. For each dimension $d$ of the flow field we can thus solve for $\tilde{\boldsymbol{\alpha}}_d^*$ using linear or ridge regression.

#### Closed-form solution for Rand-Tanh and Rand-Sig kernels

The dual form Eq. (5) still depends on the kernel function, which in our examples is parameterized by a random weight matrix $J^\psi$, which we can think of as a "quenched disorder". However, if a functional form for the kernel $k(\mathbf{x}, \mathbf{x}^*)$ is known, then the post-learning dynamics can be written without explicit reference to either $W$ or $J^\psi$, simplifying our description of the system. In certain cases we can compute the form of $k$ by assuming that $k$ depends only weakly on the specific instantiation of $J^\psi$. To do so, we take a statistical physics approach of replacing individual $k(\cdot,\cdot)$ with the expectation value $\bar{k}(\cdot,\cdot)$ computed by averaging over the quenched disorder $J^\psi$. That is,

$$k(\mathbf{x}, \mathbf{x}^*) \rightarrow \bar{k}(\mathbf{x}, \mathbf{x}^*) \equiv \mathbb{E}_{J^\psi}\left[k(\mathbf{x}, \mathbf{x}^*)\right]. \tag{24}$$

We calculate the expectation $\bar{k}$ of the kernel with respect to $J^\psi$, under the assumption that individual instantiations of the kernel approach this expectation as $N \rightarrow \infty$. For the Rand-Tanh nonlinearity, $\psi(\mathbf{x}) = \tanh(J^\psi\mathbf{x})/\sqrt{N}$, with $J_{id}^\psi \sim \mathcal{N}(0, g^2)$, this expectation is

$$\begin{aligned}
\bar{k}(\mathbf{x}, \mathbf{x}^*) = \mathbb{E}_{J^\psi}\left[k(\mathbf{x}, \mathbf{x}^*)\right] &= \frac{1}{N}\int \tanh(J^\psi\mathbf{x})^T \tanh(J^\psi\mathbf{x}^*) \prod_{i,d} P(J_{id}^\psi)dJ_{id}^\psi \\
&= \frac{1}{N}\int [\tanh(J_{1,:}\mathbf{x})\tanh(J_{1,:}\mathbf{x}^*) + \ldots \\
&\quad + \tanh(J_{N,:}\mathbf{x})\tanh(J_{N,:}\mathbf{x}^*)] \prod_{i,d} P(J_{id}^\psi)dJ_{id}^\psi \\
&= \int \tanh(J_{1,:}\mathbf{x})\tanh(J_{1,:}\mathbf{x}^*) \prod_{d=1}^{D} P(J_{1d}^\psi)dJ_{1d}^\psi
\end{aligned} \tag{25}$$

where $J_{1,:} \in \mathbb{R}^D$ and in the last line we have used the fact that each term in the sum contributes equally since all $J_{id}^\psi$ are i.i.d. For the Rand-Sig nonlinearity, $\boldsymbol{\psi}(\mathbf{x}) = (1 + \tanh(J^\psi\mathbf{x}))\sqrt{2/N}$, we arrive at a similar result:

$$\bar{k}(\mathbf{x}, \mathbf{x}^*) = 2\int \frac{1 + \tanh(J_{1,:}\mathbf{x})}{2}\frac{1 + \tanh(J_{1,:}\mathbf{x}^*)}{2} \prod_{d=1}^{D} P(J_{1d}^\psi)dJ_{1d}^\psi \tag{26}$$

The preceding two integrals can be solved exactly in the $g \rightarrow \infty$ limit. We first note that

$$\tanh(J_{1,:}\mathbf{x}) \approx \begin{cases} -1 & J_{1,:}\mathbf{x} < 0 \\ +1 & J_{1,:}\mathbf{x} > 0. \end{cases} \tag{27}$$

This means that $\tanh(J_{1,:}\mathbf{x})$ and $\tanh(J_{1,:}\mathbf{x}^*)$ are both independent of $r$ (the distance to the origin), since the norm of $J_{1,:}$ will not change the sign of the dot product, and we are in the regime where most of the probability mass is over large-norm $J_{1,:}$. This also means that the product of tanhs is either +1 or -1 (and the corresponding product n the Rand-Sig kernel evaluating to either 0 or 1), depending on whether $J_{1,:}$ has the same sign dot product with both $\mathbf{x}$ and $\mathbf{x}^*$ or different signs. If we compute the integral shell-by-shell, we realize the following. (1) The integral evaluates to $\sim 1$ when $\mathbf{x}$ and $\mathbf{x}^*$ are perfectly aligned (since the integrand becomes 1 $\times$ a probability distribution); (2) for a shell of a fixed radius $r$, the probability measure does not depend on the angle (due to the angular symmetry of products of identical Gaussians); (3) the fraction of the shell that is +1 is $(\pi - \theta(\mathbf{x}, \mathbf{x}^*))/\pi$, and the fraction of the shell that is -1 (or 0 for the Rand-Sig kernel) is $\theta(\mathbf{x}, \mathbf{x}^*)/\pi$, where $\theta(\mathbf{x}, \mathbf{x}^*)$ is the angle between $\mathbf{x}$ and $\mathbf{x}^*$.

Therefore

$$\lim_{g \to \infty} \bar{k}(\mathbf{x}, \mathbf{x}^*) = \frac{\pi - 2\arccos(\mathbf{x}^T\mathbf{x}^*/\|\mathbf{x}\|\|\mathbf{x}^*\|)}{\pi} \tag{28}$$

for the Rand-Tanh kernel and similarly

$$\lim_{g \to \infty} \bar{k}(\mathbf{x}, \mathbf{x}^*) = \frac{\pi - \arccos(\mathbf{x}^T\mathbf{x}^*/\|\mathbf{x}\|\|\mathbf{x}^*\|)}{\pi} \tag{29}$$

for the Rand-Sig kernel. Thus, in the $g, N \to \infty$ limit the learned dynamics can be expressed exactly in dual form in terms of the support states without explicit reference to either $W$ or $J^\psi$. This derivation is highly similar to that used in the literature on Neural Tangent Kernels[89].

#### Span of $1 - \theta(\mathbf{x}, \mathbf{x}^*)/\pi$ basis functions

We would like to determine the span of the functions $k_\theta(\boldsymbol{\gamma}, \boldsymbol{\gamma}^*) = 1 - \theta(\boldsymbol{\gamma}, \boldsymbol{\gamma}^*)/\pi$ for $\boldsymbol{\gamma}^*$ on the unit $(D-1)$-sphere, where $\theta(\boldsymbol{\gamma}, \boldsymbol{\gamma}^*) \equiv \arccos(\boldsymbol{\gamma}^T\boldsymbol{\gamma}^*)$. Alternatively, $k_\theta = 1/2 + \arcsin(\boldsymbol{\gamma}^T\boldsymbol{\gamma}^*)/\pi$. Our conjecture is that these functions span the set of odd functions (on the sphere) plus a constant offset, a set of functions we will call $\mathcal{G}$. We will prove this for $D = 2, 3$ but leave the statement as a conjecture for $D > 3$.

Our approach is to suppose that there is a function $h(\boldsymbol{\gamma}) \in \mathcal{G}$ of this variety that is orthogonal to all $k_\theta$, then show that it must be the zero function. Specifically, define $h \in \mathcal{G}$ such that

$$0 = \int_{S^D} h(\boldsymbol{\gamma}) k_\theta(\boldsymbol{\gamma}, \boldsymbol{\gamma}^*) d\boldsymbol{\gamma}, \tag{30}$$

where $S^D$ is the unit (D-1)-sphere (embedded in $\mathbb{R}^D$). Our task is now to show that $h = 0$. To do so, we (1) note that Eq. (30) can be rewritten as a convolution, (2) use the convolution theorem to show that all coefficients of $h$ must be zero.

For (1) observe that because $k_\theta$ depends only on the angle $\theta$, we can write

$$\int_{S^D} h(\boldsymbol{\gamma}) k_\theta(\boldsymbol{\gamma}, \boldsymbol{\gamma}^*) d\boldsymbol{\gamma} = \int_{S^D} h(\boldsymbol{\gamma}) R(\boldsymbol{\gamma}^*) k_\theta(\boldsymbol{\gamma}, 0) d\boldsymbol{\gamma} \equiv h * k_\theta \tag{31}$$

where $R(\boldsymbol{\gamma}^*)$ is a rotation operator that rotates $k_\theta(\boldsymbol{\gamma}, 0)$ by $\boldsymbol{\gamma}^*$. This is analogous to the shift operator in standard convolutions (where the argument of the final convolution is the shift amount). Note that the symmetry of $k_\theta(\boldsymbol{\gamma}, \boldsymbol{\gamma}^*)$ (i.e. it depends only on the angle between the is crucial since it means that all paths for rotating $k_\theta(\boldsymbol{\gamma}, 0)$ to a specific point $\boldsymbol{\gamma}^*$ on the sphere produce the same result.

For (2), first let $D = 2$. Then Eq. (31) is a standard convolution of a periodic function, and we can use the convolution theorem:

$$\widetilde{h * k_{\theta n}} = \tilde{h}_n \tilde{k}_{\theta n}, \tag{32}$$

where $\tilde{\cdot}_n$ is the $n$-th Fourier coefficient.

First, our definition of $\mathcal{G}$ implies that $h$ contains only a DC ($n = 0$) and odd terms (with respect to reflection of $\gamma$ about the origin). Similarly, the Fourier series of the triangle wave (given by $k_\theta$ as a function of $\theta$) has a positive DC component, and nonzero odd terms: $\tilde{k}_{\theta n} = (-1)^{(n-1)/2}/n^2$ for $n = 1, 3, 5, \ldots$. Thus, $\widetilde{h * k_{\theta n}} = \tilde{h}_n \tilde{k}_{\theta n} = 0$ implies that $\tilde{h}_n = 0$ for all $n$. Thus, $h = 0$.

For $D = 3$ we have a similar argument. By our definition of $h$, its decomposition into spherical harmonics $Y_l^m(\theta)$ will have only nonzero coefficients for $l = 0, 1, 3, 5$ and will be zero for all other even $l$. Since $k$ depends only the dot product $\boldsymbol{\gamma}^T \boldsymbol{\gamma}^*$, we can use the circular/spherical convolution theorem[24,90]:

$$F(\boldsymbol{\gamma}^*) = (h * k)(\boldsymbol{\gamma}^*) \qquad \Longrightarrow \qquad \tilde{F}_{lm} = \Lambda_l \tilde{h}_l^m \tilde{k}_l^0 \tag{33}$$

where $\Lambda_l = \sqrt{4\pi/(2l+1)}$ and $\tilde{F}_{lm}$, $\tilde{k}_{l0}$, and $\tilde{h}_{lm}$ are the $(l, m)$ coefficients of the expansion of $F$, $k_\theta$, and $h$, respectively, in spherical harmonics. (Note that $\tilde{k}_{lm} = 0$ for $m \neq 0$, a consequence of the fact that $k$ is *zonal*, depending only on the polar angle $\theta$ and not on the azimuthal angle $\phi$.)

A spherical harmonic decomposition of $k$ reveals that

$$\begin{aligned} \tilde{k}_l &= 0 \qquad l = 2, 4, 6, \ldots \\ \tilde{k}_l &> 0 \qquad l = 0, 1, 3, 5, 7, \ldots \end{aligned} \tag{34}$$

This is because, rewriting $k = .5 + k'$, where $k' = .5 - \theta/\pi$, we have that

$$\int_0^\pi k Y_l^0(\theta) \sin\theta d\theta = \int_0^\pi .5 Y_l^0(\theta) \sin\theta d\theta + \int_0^\pi k' Y_l^0(\theta) \sin\theta d\theta. \tag{35}$$

From the properties of the spherical harmonics, the first term is zero except for when $Y_0^0$, which yields a positive integral. The second term is zero for all even terms, since $Y_l^0(\theta) = Y_l^0(\pi - \theta)$ for $l$ even. For $l$ odd, however, $Y_l^0(\theta) = -Y_l^0(\pi - \theta)$, so the first term in Eq. (35) vanishes. We thus need to show that

$$\int_0^\pi k' Y_l^0(\theta) \sin\theta d\theta = \frac{2l+1}{4\pi} \int_0^\pi \left( .5 - \frac{\theta}{\pi} \right) P_l(\cos(\theta)) \sin\theta d\theta > 0, \tag{36}$$

where $P_l$ is the $l$-th Legendre polynomial. The substitution $u = \cos\theta, du = -\sin\theta d\theta$ yields

$$\int_0^\pi k' Y_l^0(\theta) \sin\theta d\theta = \frac{1}{2\pi^2} \left( \frac{2l+1}{2} \int_{-1}^1 \arcsin(u) P_l(u) du \right) \tag{37}$$

where we note that the term inside the parenthesis is the $l$-th Legendre coefficient of the arcsin function, which vanishes for $l$ even and for $l$ odd is given by $\pi/4$ for $l = 1$ and

$$= \frac{\pi}{2} \left( \left[ \frac{(l-2)!!}{((l-1)/2)! 2^{(l-1)/2}} \right]^2 - \left[ \frac{(l)!!}{((l-1)/2+1)! 2^{(l-1)/2+1}} \right]^2 \right) \tag{38}$$

for $l = 3, 5, \ldots$, which is positive (demonstrated by substituting $l = 2n + 1$).

Finally, using Eq. (34) (spherical convolution theorem for zonal filter) and the fact that $\Lambda_l \tilde{k}_l$ is positive, we must have that $\tilde{h}_{lm} = 0$ for all remaining $l = 0, 1, 3, \ldots$. Therefore $h = 0$, hence the only function in $G$ that is orthogonal to all the basis functions is the zero function, so $\{k\}$ span $G$.

##### K-WTA nonlinearity

For examples involving the K-WTA nonlinearity, we use $\psi(\mathbf{x}) = \text{K-WTA}(\tanh(J^{\psi}\mathbf{x}))/\sqrt{K}$, where K-WTA$(\mathbf{z})$ sets all except the top $K$ elements of $\mathbf{z}$ to 0.

#### Discrete-time formulation of dynamics for computation graph analysis

Here we rewrite our dynamics and PRISM plasticity equations (Eq. (1), (3)) in discrete time.

$$\mathbf{x}^t = \mathbf{x}^{t-1} + \frac{\Delta t}{\tau}\left(-\mathbf{x}^{t-1} + \mathbf{f}_{base}(\mathbf{x}^{t-1}) + C(W^{t-1})^T\boldsymbol{\psi}(\mathbf{x}^{t-1}) + B\mathbf{u}^t\right) \tag{39}$$

$$\mathbf{w}_q^t = \mathbf{w}_q^{t-1} + \Delta t I_q^t \boldsymbol{\psi}(\mathbf{x}^{t-1}) \tag{40}$$

#### E-prop implementation

The e-prop algorithm proposes[79] admits an ideal (exact) and approximate (using only purely local processes) factorization of the loss gradient.
The ideal factorization is

$$\frac{dLoss}{d\mathbf{w}_i} = \sum_t \frac{dLoss}{dz_i^t}\left[\frac{\partial z_i^t}{\partial \mathbf{w}_i}\right]_{local} = \sum_t \frac{dLoss}{dz_i^t}\mathbf{e}_i^t = \sum_t I_i^t \mathbf{e}_i^t. \tag{41}$$

The approximate factorization is

$$\frac{dLoss}{d\mathbf{w}_i} \approx \sum_t \frac{\partial Loss}{\partial z_i^t}\left[\frac{\partial z_i^t}{\partial \mathbf{w}_i}\right]_{local} = \sum_t \frac{\partial Loss}{\partial z_i^t}\mathbf{e}_i^t = \sum_t I_i^t \mathbf{e}_i^t. \tag{42}$$

We apply e-prop and BPTT to our network model using the computation graph shown in Fig S5, using an identity activation function $\mathbf{z} = \mathbf{x}$. Using either the ideal vs. approximate factorization of e-prop did not noticeably affect our results.

#### Description of the vanishing/exploding gradient problem

As described in detail in[48], the total derivative of the loss at time $T$ with respect to the parameters $\theta$ in a traditional recurrent neural network is given by

$$\frac{dLoss^T}{d\theta} = \sum_{t=1}^{T} \frac{\partial Loss^T}{\partial \mathbf{x}^T}\frac{d\mathbf{x}^T}{d\mathbf{x}^t}\frac{\partial \mathbf{x}^t}{\partial \theta} \qquad \frac{d\mathbf{x}^T}{d\mathbf{x}^t} = \prod_{t'=t+1}^{T}\frac{d\mathbf{x}^{t'}}{d\mathbf{x}^{t'-1}}. \tag{43}$$

In our setup (Fig S5), we have

$$\frac{d\mathbf{x}^{t'}}{d\mathbf{x}^{t'-1}} = \frac{\partial \mathbf{x}^{t'}}{\partial \mathbf{x}^{t'-1}} + \frac{\partial \mathbf{x}^{t'}}{\partial \boldsymbol{\psi}^{t'-1}}\frac{\partial \boldsymbol{\psi}^{t'-1}}{\partial \mathbf{x}^{t'-1}} = (I - \frac{1}{\tau}) + CW^T\text{diag}(\sigma')J^{\psi} \tag{44}$$

where diag$(\sigma')$ is a diagonal matrix of the derivatives of the nonlinearity, e.g. Sigmoid or Tanh. Absent strong constraints on $W$ and $J^{\psi}$, the product of many such terms will either vanish or explode when used to calculate gradients in BPTT.

#### Dual form of the dynamics with temporally extended plasticity windows

Consider an arbitrary temporal plasticity kernel, which specifies the weight update given delta pulses in the instructive signal and presynaptic input as a function of their time delay $\Delta t_{iq} = t_i - t_q$:

$$\Delta W_{iq}^{\delta} = h(\Delta t_{iq}). \tag{45}$$

The corresponding dynamics for the weight updates are then

$$\frac{dW_{iq}}{dt} = (I_q * h)(t)\psi_i(\mathbf{x}(t)) \tag{46}$$

or equivalently

$$\frac{dW_{iq}}{dt} = I_q(t)(h_- * \psi_i)(t) \tag{47}$$

where $h_-(\Delta t) \equiv h(-\Delta t)$. Note that the above reduces to the simplified learning rule (Eq. (3)) when $h(\Delta t_{iq}) = \delta(\Delta t_{iq})$. When the instructive signal is a series of delta pulses

$$\mathbf{I}(t) = \sum_l \boldsymbol{\alpha}_l^* \delta(t - t_l^*) \tag{48}$$

the time integral of the weight update rule reduces to

$$\int \frac{dW_i}{dt} dt = \int_t \sum_l \boldsymbol{\alpha}_l^* \delta(t - t_l^*)(h_- * \psi_i)(t)dt' = \sum_l \boldsymbol{\alpha}_l^* (h_- * \psi_i)(t_l^*) \equiv \sum_l \boldsymbol{\alpha}_l^* \tilde{\psi}_i^l, \tag{49}$$

yielding

$$\Delta W_{i,:} = \sum_l \boldsymbol{\alpha}_l \int_{t'} h(t')\psi_i(\mathbf{x}(t_l^* - t'))dt' \tag{50}$$

such that

$$\Delta \mathbf{f} = C \sum_l \int_{t'} h(t')\boldsymbol{\psi}(\mathbf{x}(t_l^* - t'))^T \boldsymbol{\psi}(\mathbf{x})dt' = C \sum_l \int_{t'} h(t')k(\mathbf{x}, \mathbf{x}(t_l^* - t'))dt'. \tag{51}$$

A visualization of this result is shown in Fig S6.

#### Details of the case studies in the *online learning of unknown dynamics real-time error signals* section

In both of the case studies analyzed (Fig 4), the *reference system* is a Van der Pol oscillator, a particular case of Liénard system[55], with dynamics:

$$\ddot{z}(t) + \mu(z(t)^2 - 1)\dot{z}(t) + z(t) + u(t) = 0, \tag{52}$$

where $\mu > 0$ is a parameter. We use $\mu = 2$ in our case studies. We define the reference system's state as

$$\mathbf{z}(t) := \begin{bmatrix} z(t) \\ \dot{z}(t) \end{bmatrix}, \tag{53a}$$

and accordingly the state-space representation of the dynamics (52) is:

$$\dot{\mathbf{z}}(t) = \underbrace{\begin{bmatrix} \dot{z}(t) \\ -\mu\big(z(t)^2 - 1\big)\dot{z}(t) - z(t) \end{bmatrix}}_{=:\mathbf{f}_{ref}(\mathbf{z})} + \underbrace{\begin{bmatrix} 0 \\ -1 \end{bmatrix}}_{=:B} u(t). \tag{53b}$$

The only attractor in the autonomous Van der Pol oscillator is a stable limit cycle, which yields self-sustained relaxation oscillations for large $\mu$ values. The origin is an unstable fixed point. We consider two case studies, corresponding to different RNN base dynamics. We set $\tau = 1$.

- **Case study 1:** $\mathbf{f}_{base} \equiv \mathbf{0}$ and thus, when $W = 0$ the RNN behaves as a stable linear system with exponential convergence rate to the origin (see Fig 4b, Case 1, 'before learning'). The RNN follows the dynamics

$$\frac{d\mathbf{x}}{dt} = -\mathbf{x} + CW^\top \boldsymbol{\psi}(\mathbf{x}) + \kappa \mathbf{e} + B\mathbf{u}. \tag{54}$$

- **Case study 2:** the superposition of the leakage term and base dynamics yields a bi-stable attractor, with stable fixed points $\mathbf{x}_\pm = [\pm 1 \ \pm 1]^\top$ (see Fig 4b, Case 2, 'before learning'). The RNN follows the dynamics

$$\frac{d\mathbf{x}}{dt} = -\frac{1}{8}(A_1\mathbf{x})^3 + A_2\mathbf{x} + CW^\top \boldsymbol{\psi}(\mathbf{x}) + \kappa \mathbf{e} + B\mathbf{u}, \tag{55}$$

  where $A_1 = \begin{bmatrix} 1 & 1 \\ 1 & 1 \end{bmatrix}$ and $A_2 = \begin{bmatrix} 0 & 1 \\ 1 & 0 \end{bmatrix}$.

In both case studies we set the entries $C_{dq} \sim \mathcal{N}(0, 1)$ i.i.d. ($d = 1, \ldots, D$ and $q = 1, \ldots, Q$) with $D = 2$ and $Q = 10$. The control gain is set to $\kappa = 5$ and $N = 75$ with the Rand-Tanh kernel $\psi(\mathbf{x}) = \tanh(J^\psi \mathbf{x})/\sqrt{N}$, with $J^\psi_{id} \sim \mathcal{N}(0, 1)$ i.i.d. ($i = 1, \ldots, N$). The learning rate is set at $\eta = 1$. During the instructed learning period, $\mathbf{u}$ is white Gaussian noise.

### SUPPLEMENTAL FIGURES

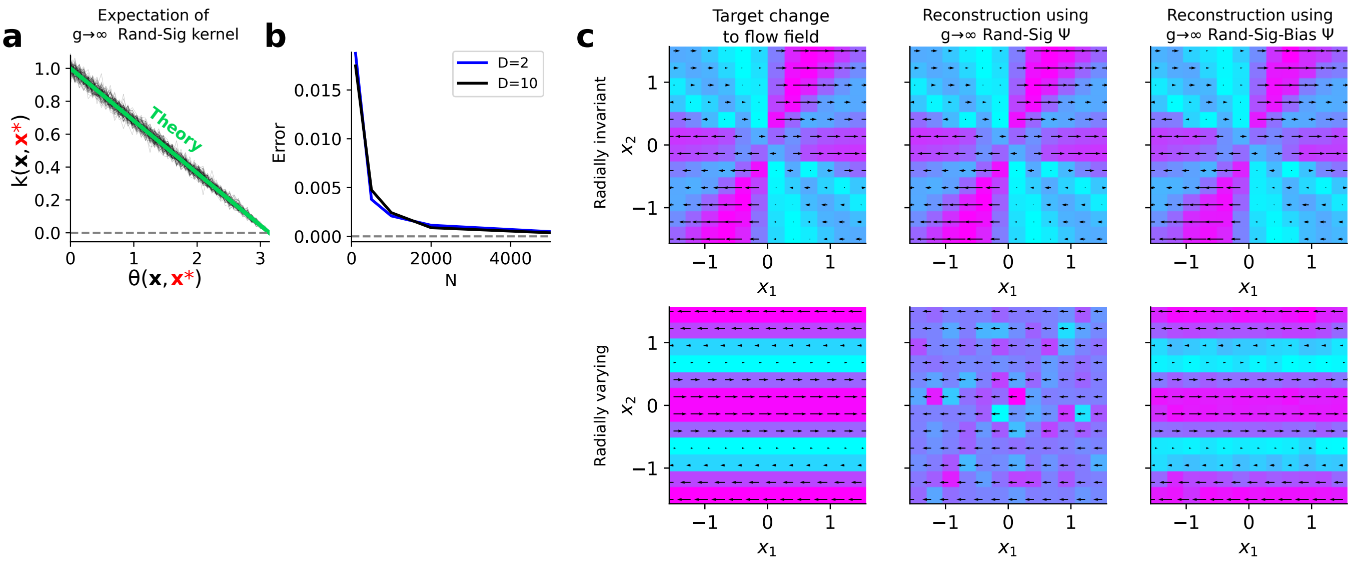


**Figure S1: Example flow field changes achievable from support states**

**a**. Theoretical prediction vs simulation of kernel value for two support states for using the Rand-Sig-$\infty$ kernel, as a function of the angular distance between the support states. **b**. Root-mean-squared error between theoretical and numerical prediction of kernel value as a function of $N$, for two different $D$. **c**. Example target flow field changes with (top) and without (bottom) radial invariance, and their reconstructions ($\{\alpha_l^*\}$ identified through ridge regression) using two different $\psi(\cdot)$, $k(\cdot,\cdot)$, each using 100 support states with uniformly distributed angles and random radii. The Rand-Sig-Bias nonlinearity is $\psi(\mathbf{x}) = (1 + \tanh(J^\psi \mathbf{x} + \mathbf{b}))\sqrt{2/N}$, where $b_i \sim \mathcal{N}(0, g)$.

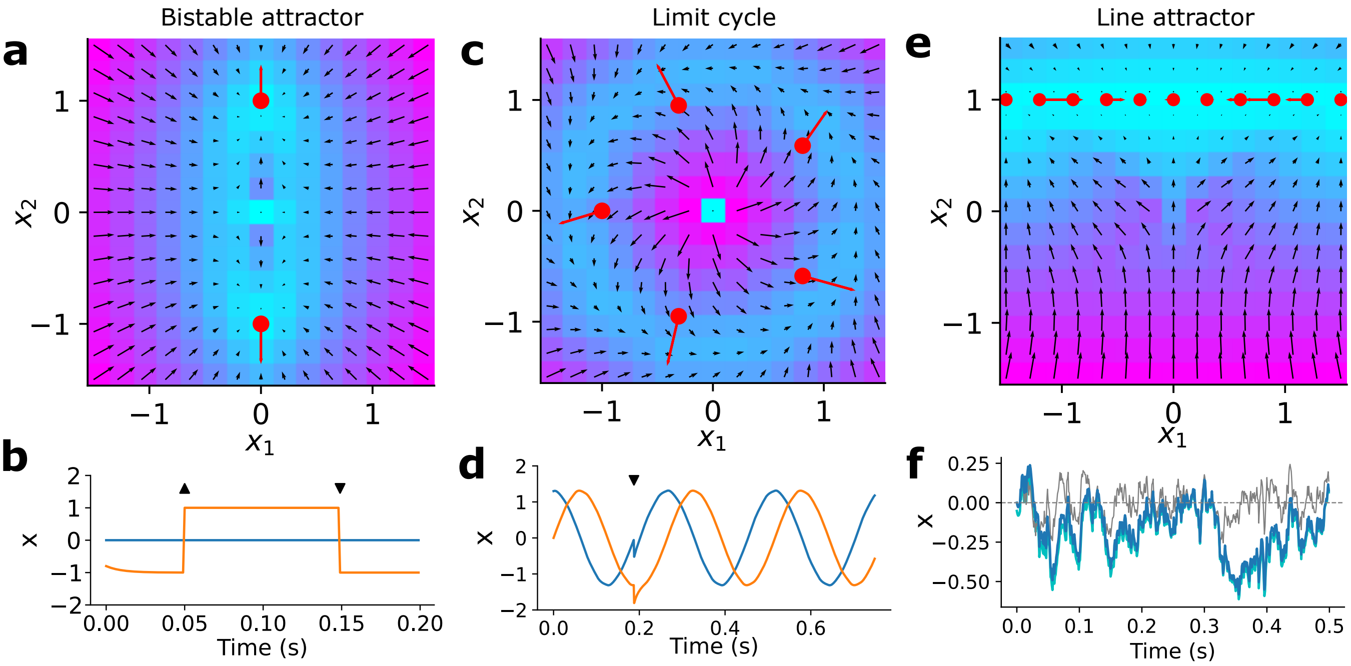


**Figure S2: ridge regression construction of attractor networks**

**a**. Solution to flow field producing bistable attractor. Red dots indicate support states and lines directions of associated feedback forces. **b**. Example time-course produced via the flow field in **a**. Upward arrow indicates positive delta-pulse input to $x_2$; downward arrow indicates negative delta-pulse input to $x_2$. Blue: $x_1$, orange: $x_2$. **c.** As in **a** but for a limit cycle. **d.** Trajectory produced via flow field in **c**. Downward triangle indicates negative perturbation to both $x_1$ and $x_2$, demonstrating the stability of the limit cycle. **e.** As in **a** but for a line attractor. **f.** Timecourse of $x_1$ (dark blue) in response to a white noise input along $x_1$ direction. A constant unity input is provided in the $x_2$ direction (such that the line attractor exists along $x_2 = 1$). Cyan indicates true time-integral of the input. Gray line indicates $x_1$ response before support states are added.

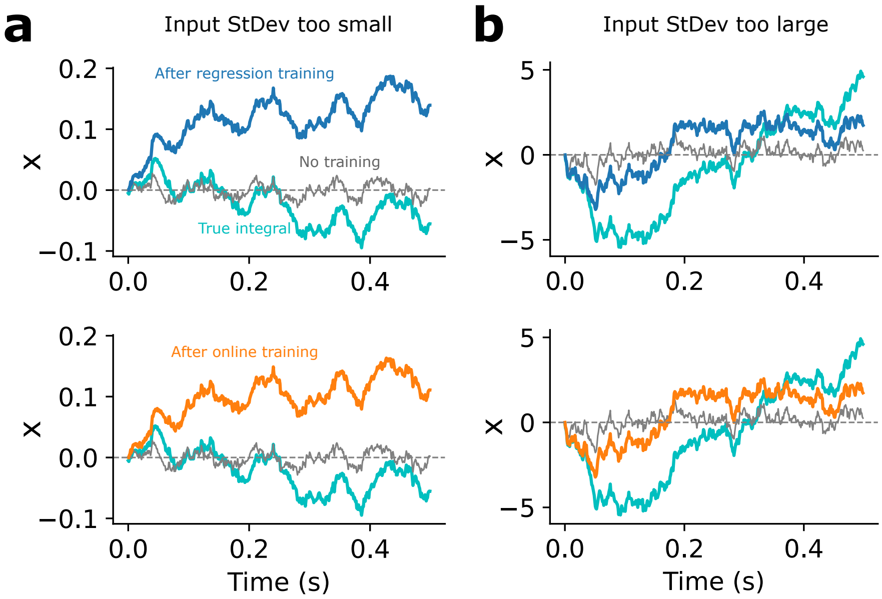


**Figure S3: Line attractor failure modes**

**a.** Response of the approximate line attractor in Fig 3g to a white noise input with a small standard deviation, when the flow field is created using either ridge regression (top) or online (bottom). **b.** As in **a** but when the input standard deviation is too large.

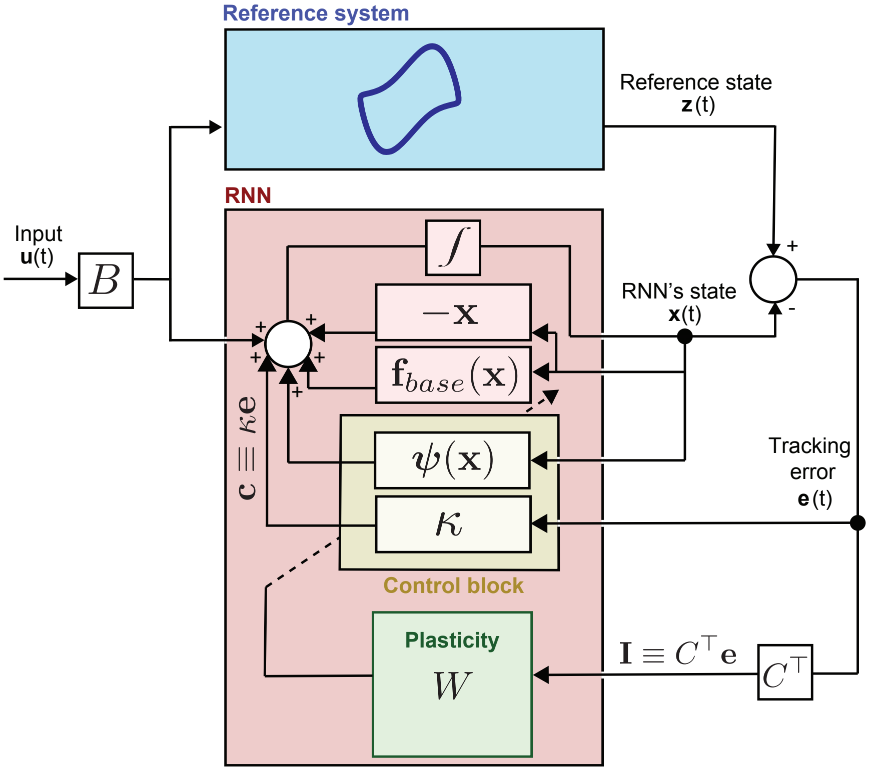

**Figure S4: Block diagram for online learning of unknown dynamics**
The external input $\mathbf{u}$ enters both the reference system (blue) and the RNN (red). The tracking error signal $\mathbf{e}$ defined as the difference between the reference system's and RNN's states (i.e., $\mathbf{e} \equiv \mathbf{z} - \mathbf{x}$) is fed back into the RNN to produce the instructive ($\mathbf{I} \equiv C^\top \mathbf{e}$) and corrective ($\mathbf{c} \equiv \kappa \mathbf{e}$) signals responsible for plasticity, as indicated in the diagram. The adaptive control block (yellow) can be interpreted as the composition of a nominal component (the corrective signal $\mathbf{c}$) and an adaptive component (the signal impacted by plasticity, $C^\top W \psi(\mathbf{x})$). The dotted arrow from the plasticity block to the control block indicates adaptation. We remark that the flow field of the reference model is *not* known by the RNN — only the tracking error signal $\mathbf{e}$ is observed by the RNN during instructed learning in a streaming (online) fashion.

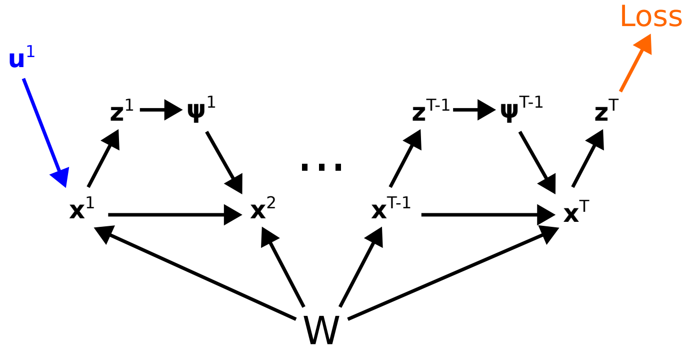


**Figure S5: Hold task computation graph for BPTT and e-prop**
Depiction of the computation graph used in the hold task in Fig 5 for BPTT and e-prop, in which the loss is evaluated only after a delay relative to the input. Note that the weights $W$ are static over the forward pass. To implement e-prop[79] we introduce an explicit variable $\mathbf{z}^t \in \mathbb{R}^D$, corresponding to the "activation" of the neurons $\phi(\mathbf{x}^t)$. Although we use the identity activation $\phi(\mathbf{x}^t) = \mathbf{x}^t$, which does not change the dynamics, this is necessary for the e-prop factorization.

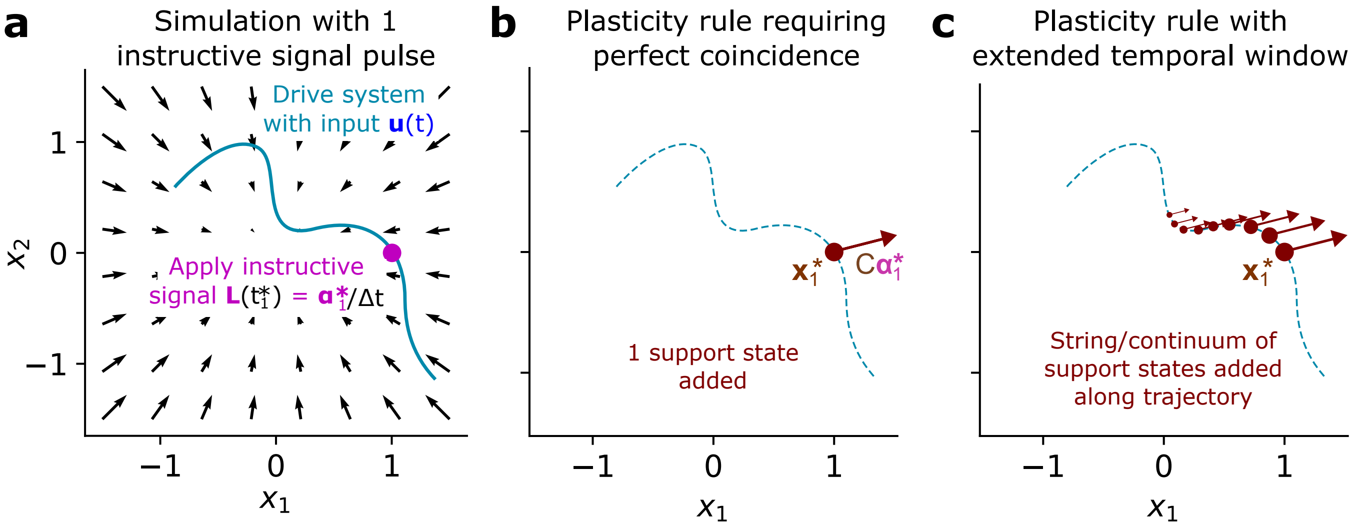


**Figure S6: Dual form for temporally extended plasticity window**
**a**. Simulation schematic in which system is driven input and a single instructive signal pulse is applied at $t_1^*$. (As in Fig 2a.) **b**. Result of instructive signal in **a** using plasticity rule with perfect coincidence (Eq. (3)). **c**. Schematic of result of instructive signal in **a** using plasticity rule with a temporally extended plasticity window, in which coincidence is defined by a decaying temporal kernel between the instructive signal and presynaptic activity.

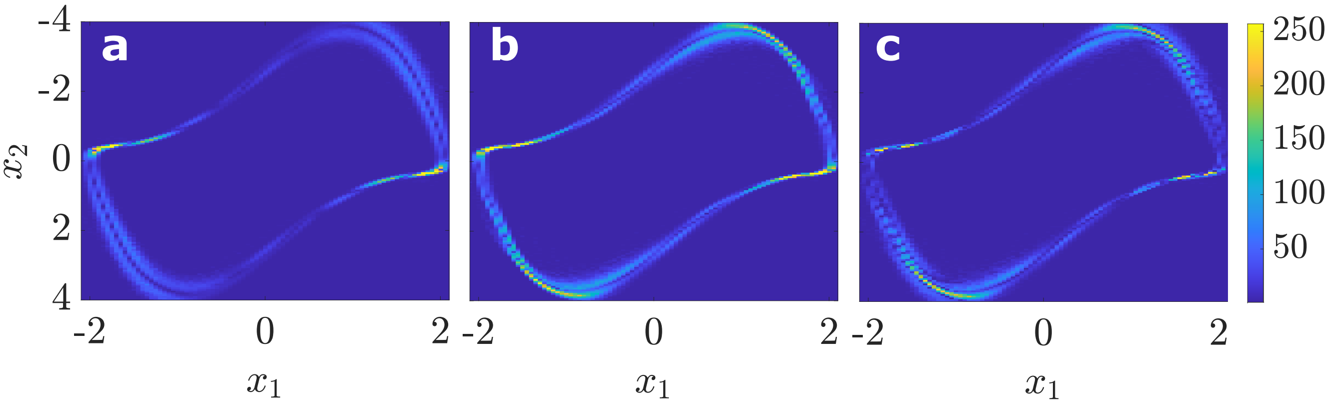


**Figure S7: Support state density of continuous learning signals**
Norm of the density $\boldsymbol{\rho}$, as defined in Eq. (19), for the continuous learning signal method developed in the section *Online learning of unknown reference dynamics through adaptive control*. The color code is according to the norm of the cumulative value of alpha along the training trajectories: $\| \int_0^T \boldsymbol{\alpha}(t)dt \|$, where $T$ denotes the learning horizon. Colorcode is consistent among panels, as given by the colorbar, and so is the vertical axis. **a.** Case 1: base field is composed by the leakage term. **b.** Case 2: base field is the bi-stable attractor. **c.** Absolute value of the difference in $\| \int_0^T \boldsymbol{\alpha}(t)dt \|$ between Case 1 and Case 2. The case studies and parameter values are given in Fig 4 and in Extended Methods.
